# Coding for Circuit Integration in the Injured Brain by Transplanted Human Neurons

**DOI:** 10.1101/2025.08.14.670350

**Authors:** Zhifu Wang, Danyi Zheng, Shu-Min Chou, Phil Jun Kang, Fei Ye, Su-Chun Zhang

**Affiliations:** Program in Neuroscience and Behavioral Disorders, Duke-NUS Medical School, Singapore; Center for Neurologic Diseases, Sanford Burnham Prebys Medical Discovery Institute, La Jolla, CA, USA

**Author notes:** These authors contributed equally (Zhifu Wang, Danyi Zheng). Senior author (Su-Chun Zhang).

## Abstract

Neural transplantation holds the potential to repair damaged neural circuits in neurological diseases. However, it remains unknown how the grafted neurons project axons to and make functional connections with the appropriate targets to repair the damaged circuit at the adult stage. Here we report that human cortical progenitors, transplanted into the ischemic mouse motor cortex, matured and integrated into cortical and subcortical neural circuits including the corticospinal tract. Neuronal tracing combined with single-nuclei RNA sequencing revealed the close relationship between the transcription profiles of a cortical neuronal subtype, especially those of axon guidance and synapse assembly, with the specific target projection and synapse organization. Machine learning-based regression further identified the transcriptional codes for the targeted projection and circuit integration to reconstruct the damaged circuits. Our finding opens a promising strategy for treating neurological diseases through promoting regeneration and neural transplantation.

**Highlights:** Human ESC-derived cortical neurons reconstitute the ischemic motor cortex Functionally repaired corticospinal tract restores animal behaviors

Transplanted cortical neurons exhibit subtype-specific projection and integration Unique transcriptional coding for axonal navigation in the mature CNS

## INTRODUCTION

The human brain has a limited regenerative capacity. Transplantation of neural progenitor cells (NPCs) offers a promising approach to replace the lost neurons and reconnect the damaged circuits. Indeed, transplantation of NPCs derived from pluripotent stem cells, including human embryonic stem cells (hESCs) or induced pluripotent stem cells (hiPSCs), is now in clinical trials for Parkinson’s disease (PD), epilepsy, and age-related macular degeneration. However, how the grafted neurons find and make functional connections with their targets in the mature injured brain remains unclear. This makes neural transplantation therapy less predictable, especially for diseases of the cerebral cortex that affect diverse neuronal subtypes and multiple neural circuits including the long-distance corticospinal tract (CST).

Cortical neurons or organoids transplanted into the postnatal brain exhibit low efficiency in extending axons from the neocortex to the brainstem^1^, with even fewer projections observed when grafted into the brains of adult animals following traumatic brain injury or stroke^2,3^. Meanwhile, the injury leaves an open cavity surrounded by glial scars, resulting in a hostile microenvironment that severely limits the survival and integration of transplanted neurons, preventing effective structural repair and reconstruction of the damaged neural circuitry. Single cell profiling of retrogradely labelled neurons revealed the self-organized process during development based on the association between the transcriptome of cortical neuronal subtypes and their projection targets^4–6^. However, the injured brain environment at adult stage differs significantly from the developing brain in both structure and molecular composition, and circuit integration relies on a complex combination of guidance molecules^7^. Substantial numbers of graft-derived axons are often observed in stochastic, unrelated brain regions. Even the neurons originating from the visual cortex, transplanted into the visual cortex, project to the agranular insula, striatum, amygdala, and superior colliculus^8–10^, only partly corresponding to the host neural circuits. On the other hand, hESC-derived midbrain dopaminergic (mDA) neurons, transplanted into the substantia nigra (SN), primarily grow axons along the medial forebrain bundle (MFB) to innervate the striatum^11,12^, whereas cortical neurons transplanted into the SN project to the prefrontal cortex^11^. These phenomena suggest that the organization principle of the projection and connectivity in the damaged adult brain is different from the self-organizing processes observed during brain development^13^. Hence, identifying the basic principle of pathfinding and circuit integration by grafted neurons in the mature brain will enable us to develop not only a predictable as well as safe and effective cell transplantation therapy but also regenerative therapy for brain disorders.

We reconstructed subcortical neuronal circuits, including the CST, in the ischemic stroke brain by transplanting subtype-specific deep-layer cortical neurons. These specific cortical neurons facilitated more efficient establishment of targeted projections and were sufficient to reconstruct long-distance neural circuits without overgrowth. This finding suggests a potential link between axonal projection directions and neuronal subtype identity. Single-nucleus RNA sequencing (snRNAseq), in combination with barcoded retrograde tracing, revealed that transplanted cortical neurons showed subtype-specific projection and integration. The subtypes of hPSC-derived cortical neurons, transplanted in the adult brain, possess distinct combinations of axon guidance and synaptic assembly molecules that are similar to but distinguishable from those required in endogenous human neurons. We further identified the transcriptional codes for specific projection and circuit integration in the mature brain with machine learning. These findings highlight the necessity of developing appropriate neural types for integration into target neural circuits in the injured brain by neural transplantation at the adult stage.

## RESULTS

### Transplanted human cortical neurons project to specific brain regions

The stroke cavity presents a hostile environment for transplanted cells; hence, grafting is typically performed in adjacent healthy brain regions instead^3,14–16^, leaving the injured cortex open and unrepaired. We previously developed a cocktail that supports the survival of neural cells transplanted into the ischemic cavity^17^. To reconstitute the ischemic cortex and restore damaged circuits, we first differentiated hESCs into cortical NPCs for transplantation (Figure 1A). The cortical NPCs at day-35 contained over 70% progenitors expressing both CTIP2 and FEZF2 (markers of layer 5 pyramidal neurons) with less than 20% expressing layer-6 marker TBR1 (Figures 1B and S1A-S1B). The ischemic stroke models were induced by photothrombosis in the motor cortex of 12-week-old immunodeficient mice (Figure S1). The cortical NPCs were transplanted into the lesion cavity of ischemic stroke models (Figure S1). In total, we successfully performed cortical neuron transplants in 95 mice (Supplementary Table 1). Twelve months post transplantation, the graft filled the stroke cavity with no overgrowth (Figures 1C and S1C).

**Figure 1.**
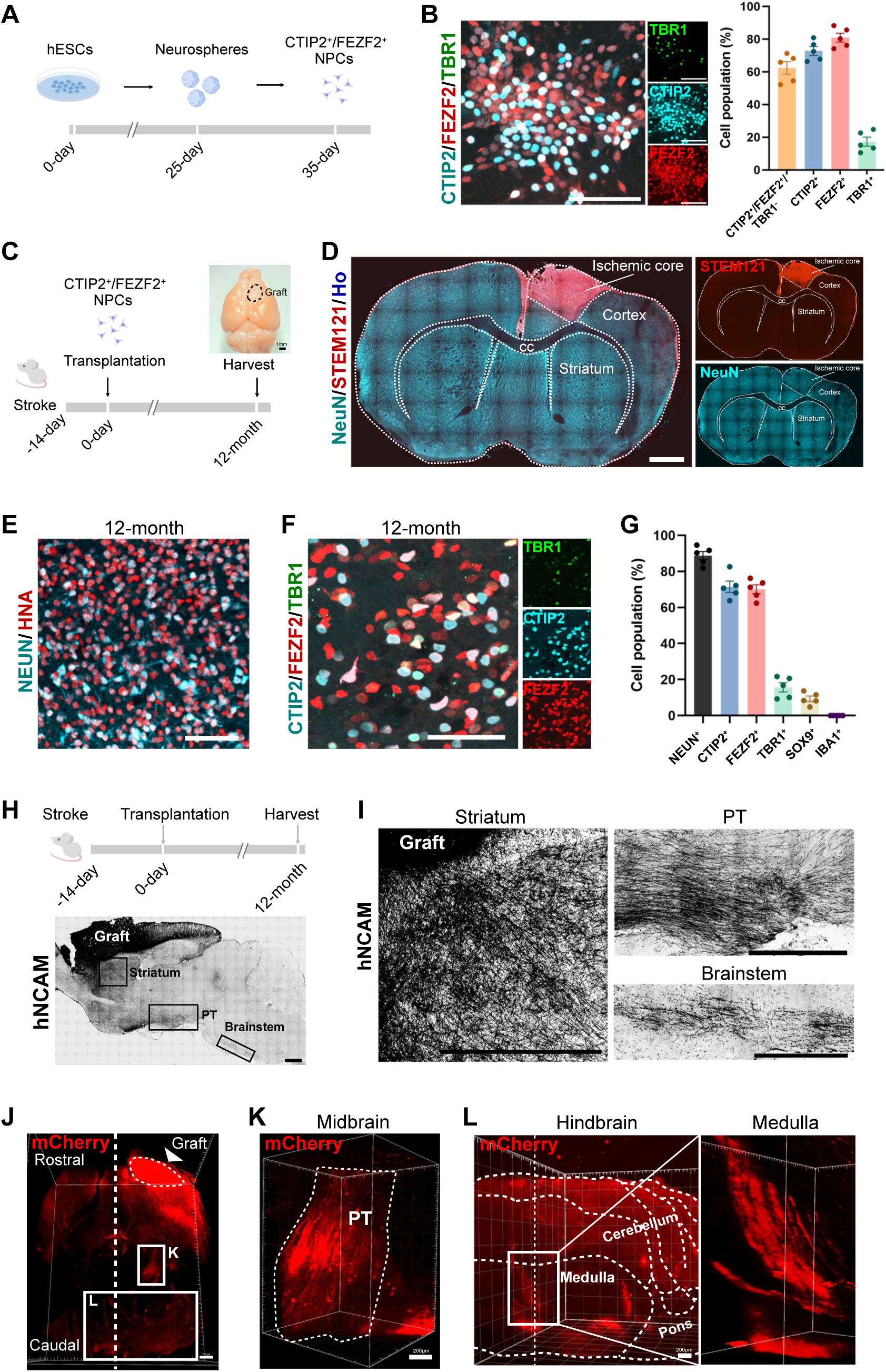
Transplanted human cortical neurons reconstitute the damaged cortex and project axons to subcortical regions. (A) Cartoon outlining the differentiation of the human cortical NPCs from hESCs. (B) Immunostaining for CTIP2, FEZF2, and TBR1 showing the cell population of induced human cortical NPCs. Quantification on the right panel; n=5 independent samples. Scale bars, 200μm. (C) Experimental strategy for transplantation of human cortical NPCs in stroke mice. The samples were collected at 12 months after transplantation. Scale bars, 1mm. (D) Immunostaining with NeuN and STEM121(human cell marker) shows that the transplanted NPCs became NeuN+ neurons and filled the ischemic cavity at 12 months post-transplantation. Scale bars, 1mm. (E) Immunostaining shows co-labelling of NEUN and human nuclear antigen (HNA) in transplanted neurons at 12 months post-transplantation. Scale bars, 200μm. (F) Immunostaining for CTIP2, FEZF2, and TBR1 on transplanted neurons at 12 months post-transplantation. Scale bars, 100μm. (G) Quantification for (F) shows the cell population of transplanted neurons at 12 months post-transplantation. n=5 mice. (H) Immunostaining for hNCAM showing axonal projections of transplanted neurons in mature mouse brains. (I) Magnification of (H) showing the axon projection by grafted neurons in the striatum, pyramidal tract (PT) and brainstem. (J to L) 3D reconstructed images of mCherry showing axon bundle derived from grafts neurons in the transverse plane (J), midbrain (K), and hindbrain (L). Scale bars, 700μm in (J), 200μm in (K) and (L). All Data are represented as mean±SEM.

Immunohistochemical analysis showed that 90% of transplanted cells expressed the neuronal marker NeuN (Figures 1D and 1E), indicating differentiation into neurons. About 63% of grafted cells were CTIP2^+^/ FEZF2^+^/ TBR1^-^ neurons (Figures 1F and 1G), consistent with the pre-transplantation cell population, suggesting that these cortical NPCs were cell fate-committed before transplantation. Single-nucleus RNA sequencing (snRNA-seq) revealed that the grafted human cells were predominantly deep layer neurons, with only a small fraction of upper layer and GABAergic neurons (Figures S1D-1H). Less than 10% of transplanted cells differentiated into astrocytes (SOX9^+^), and none into microglia and oligodendrocytes (Figures S1I-S1K). At this stage, no KI67-expressing human cells were detected (Figure S1L). To further test the sustained stability of this cell fate, we collected snRNA-seq data at pre-transplantation, three months post-transplantation, and nine months post-transplantation, and integrated them with data obtained at twelve months after transplantation (Figure S2). This analysis showed, at the single-cell transcriptomic level, the survival, proliferation, and eventual maturation of the transplanted cells over the twelve-month period (Figure S2). Thus, transplanted human NPCs survived and differentiated into deep-layer cortical neurons, with stable proportions of CTIP2⁺/FEZF2⁺ cells pre– and post-transplantation. We also observed that graft-derived axons (hNCAM^+^) were enveloped by MBP^+^ cells (Figure S3A). Since no human cells become oligodendrocytes, indicated by histology (Figure S3B) and snRNA-seq (above), the MBP^+^ cells are originated from mouse oligodendrocytes. Additionally, mouse endothelial cells (Laminin^+^) infiltrated the graft (Figure S3C), enabling blood vessels to support the grafts in a long term.

We evaluated the extension of graft-derived axons by human-specific hNCAM staining (Figure 1H), which revealed extensive axonal projections from the graft to cortical and subcortical brain regions. Semi-quantification of hNCAM^+^ axonal density in the whole brain revealed that the graft-derived axons were not only present along CST, but also in the somatosensory cortex (S1), thalamus, superior colliculus, medial vestibular nucleus, and SC (Figures S4A-S4E and S4H). In contrast, only a small number of axons followed the corpus callosum (cc) to reach the contralateral hemisphere (Figures S4F and S4H). Therefore, we focused our analysis on the projection and connectivity in the ipsilateral hemisphere. Robust projections were observed along the host CST from the striatum to the brainstem, as revealed in sagittal sections (Figure 1I). To clearly visualize graft-derived axons in the pyramidal tract, we performed tissue clearing and 3D reconstruction for mCherry^+^ transplanted neurons in the brain by light-sheet microscopy (Figure 1J). Fascicular mCherry^+^ axons extended through the midbrain along the pyramidal tract, decussated at the medulla, and continued into the spinal cord (Figures 1J-1L and S4G). Such a projection pattern of transplanted neurons (CTIP2^+^/ FEZF2^+^/ TBR1^-^) resembles that of endogenous corticospinal neurons.

### Transplanted neurons synapse with host subcortical neurons with specificity

Functional restoration depends on integration of transplanted neurons into the existing circuits. To evaluate the global post-synaptic connectivity of transplanted neurons in the adult brain and spinal cord, we developed an ESC line that expresses the anterograde trans-synaptic protein mWGA-mCherry (mWmC), a mono-synaptic tracer^18^, driven by the synapsin1 promoter (Figure 2A). In vitro, the mWmC was only detected in neurons (NeuN^+^) but not in non-neural cells (Figure S5A). We first co-cultured mWmC/GFP neural progenitor cells with wild-type neural progenitors and did not observe punctate red signal in the wild-type cells (Figure S5B), indicating that mWmC is not transmitted between progenitor cells. To further establish that mWmC functions as a monosynaptic tracer, we cultured upstream mWmC/GFP-expressing neurons and downstream hM3Dq-mCherry– labeled neurons in adjacent chambers, with mCherry labeling the cell membrane to demarcate cell body (Figure S5C). Following axonal projections from the mWmC/GFP neurons to the downstream neurons, punctate red signals were detected exclusively in the first-order postsynaptic neurons but not in second-order neurons (Figure S5C). Following transplantation, we detected mWmC signals in the host neurons, suggesting that transplanted neurons formed synaptic connections with downstream host neurons (Figure 2B). A substantial number of mWmC-positive host neurons were observed in the S1, striatum, thalamus, subthalamic nucleus (STN), superior colliculus (SuC), pontine gray (Pon G) (Figures 2C-2G). In contrast, despite extensive graft-derived axons, there were few mWmC-labelled cells observed in the substantia nigra (Figure 2G). These results indicate that transplanted neurons make connections with neurons in specific target brain regions until their axons reach the targets rather than form stochastic connections with nearby host neurons based on proximity.

**Figure 2.**
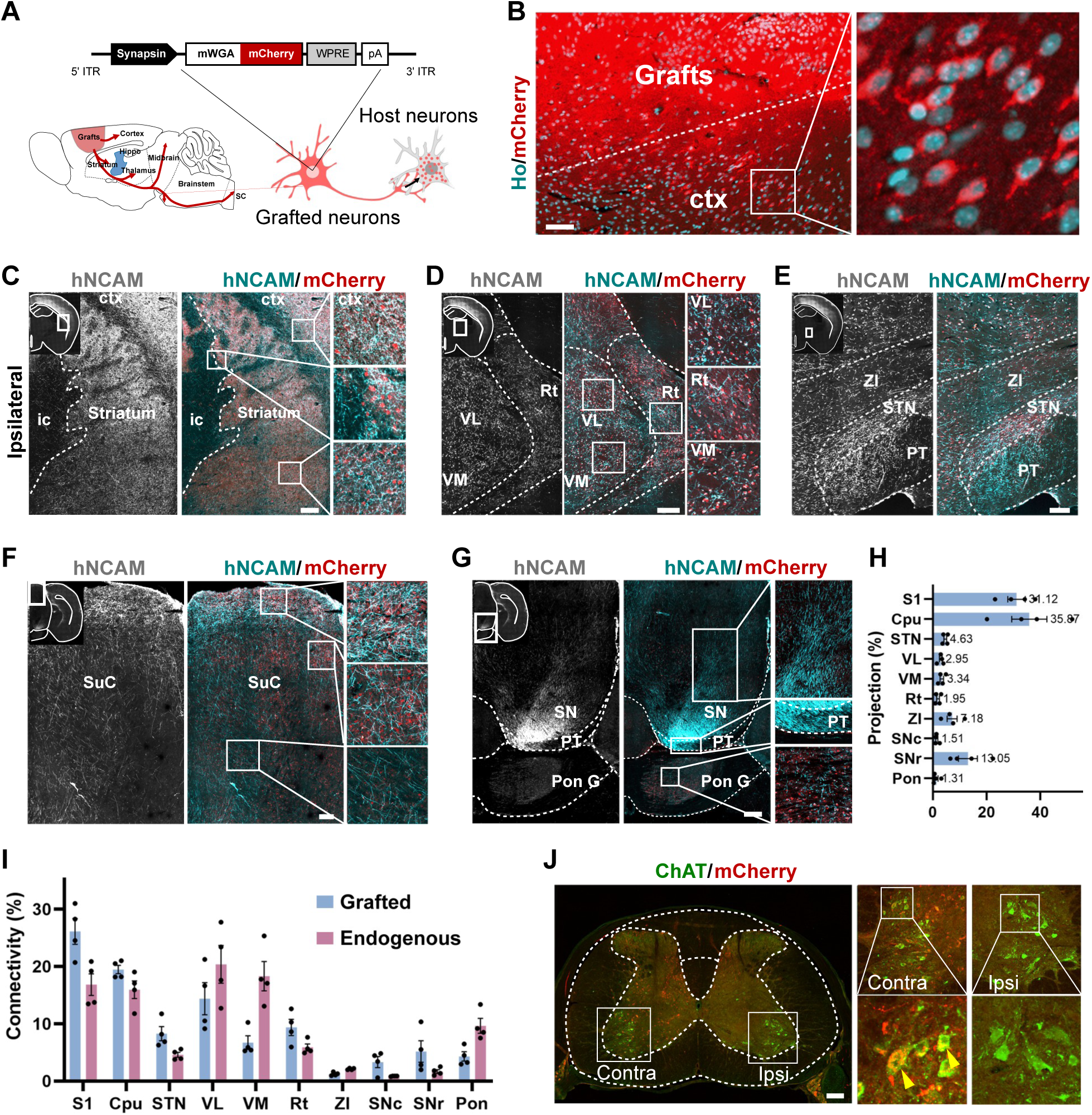
Post-synaptic connectivity is mapped by anterograde tracing. (A) Cartoon illustrating the generation of the hSyn-mWmC hESC line for anterograde tracing in the transplanted brain. (B) Immunostaining for mCherry showed the presence of mWmC in host neurons in the intact cortex adjacent to the graft. Scale bars, 100μm. (C to G) Immunostaining for hNCAM and mCherry showing axons of grafted neurons (hNCAM+) and mCherry labeled host neurons in ipsilateral S1 and striatum (C), thalamus (D), ZI and STN (E), SuC (F), Pon G and SN (G). S1, primary somatosensory cortex. ic, internal capsule. VL, ventrolateral thalamic nucleus. Rt, reticular thalamic nucleus. ZI, zona incerta. STN, subthalamic nucleus. PT, pyramidal tract. SuC, superior colliculus. SN, substantia nigra. Pon G, pontine grey. Scale bars, 100μm. (H) Quantification of axon projection by grafted cells in mature brains. n=4-5 mice. Data are represented as mean±SEM. (I) Quantification of output connections of transplanted neurons (blue) and endogenous corticospinal motor neurons (purple). The connectivity ratio is shown for all brain regions receiving synaptic input from transplanted neurons at 12-month, compared to endogenous neurons in intact M1(see Figure 7B). n=4 mice per group. Data are represented as mean±SEM. (J) Immunostaining showing WGA-labeled, ChAT-expressing host neurons in the spinal cord. Dot plot of top marker genes. Insets are magnified on the right.

We next mapped the connections of graft-derived axons, showing that the transplanted neurons projected to and established synaptic connectivity in motor-related brain areas including S1, striatum, thalamus, STN, SuC, and Pon G (Figure 2H and 2I). This is strikingly similar to the endogenous targets of the motor cortex when we mapped the post-synaptic connectivity by AAV-mWmC injection into the motor cortex (Figure S6A). Notably, mWmC-positive cells were detected in the SC, primarily on the contralateral side (Figure 2J), implying that the transplanted neurons are integrated in the corticospinal circuits. Remarkably, these transplanted human neurons connected with lower motor neurons (ChAT^+^) in the ventral horn of the spinal cord (Figure 2J), whereas rodent upper motor neurons usually form indirect connections with lower motor neurons through interneurons rather than direct connection^19–21^. To verify this distinction, we examined lower motor neurons in the spinal cord following AAV-mWmC injection in intact motor cortex and found no detectable mWmC signals in spinal motor neurons (Figure S6B), supporting the absence of direct connections rodents.

At the cellular level, our immunohistochemistry results showed that the transplanted neurons established connections with CTIP2+ spiny GABAergic projection neurons in the striatum (Figure S7A), (vGLUT1^+^) glutamatergic and (GAD67^+^) GABAergic neurons in the thalamus, and glutamatergic neurons in the STN, as indicated by the mCherry signal in those cells (Figures S7B, S7C, and S7F). Meanwhile, mWmC was not detected in dopaminergic neurons in the substantia nigra (SN) (Figure S7D). The grafted neurons formed post-synaptic connections with interneurons in the SC (Figures S7E and S7F).

### Transplanted cortical neurons make functional connections with host neurons

To test if the connections between transplanted neurons and host neurons are functional, we generated a DREADD/mWmC hESC line which expresses the excitatory hM3Dq receptor regulated by Deschloroclozapine (DCZ) and the trans-synaptic tracer mWmC (Figures 3A and S8A). This strategy allows us to simultaneously assess trans-synaptic connections using mWmC labelling and functional synaptic connectivity via DREADD activation. Administration of DCZ (0.05mg/kg) significantly increased the expression of the activity-dependent protein FOS in transplanted neurons, indicating that DCZ treatment activated the transplanted DREADD/mWmC neurons (Figure 3B). Immunohistochemical co-staining of FOS and WGA showed that following DCZ activation of the transplanted neurons, the proportion of synaptically connected host neurons (WGA^+^) expressing FOS also significantly increased, while there was no significant change in host neurons (WGA^-^) without synaptic connections with the graft (Figure 3B and 3C). These data indicate that transplanted neurons form synaptic connections with host neurons and influence their activity.

**Figure 3.**
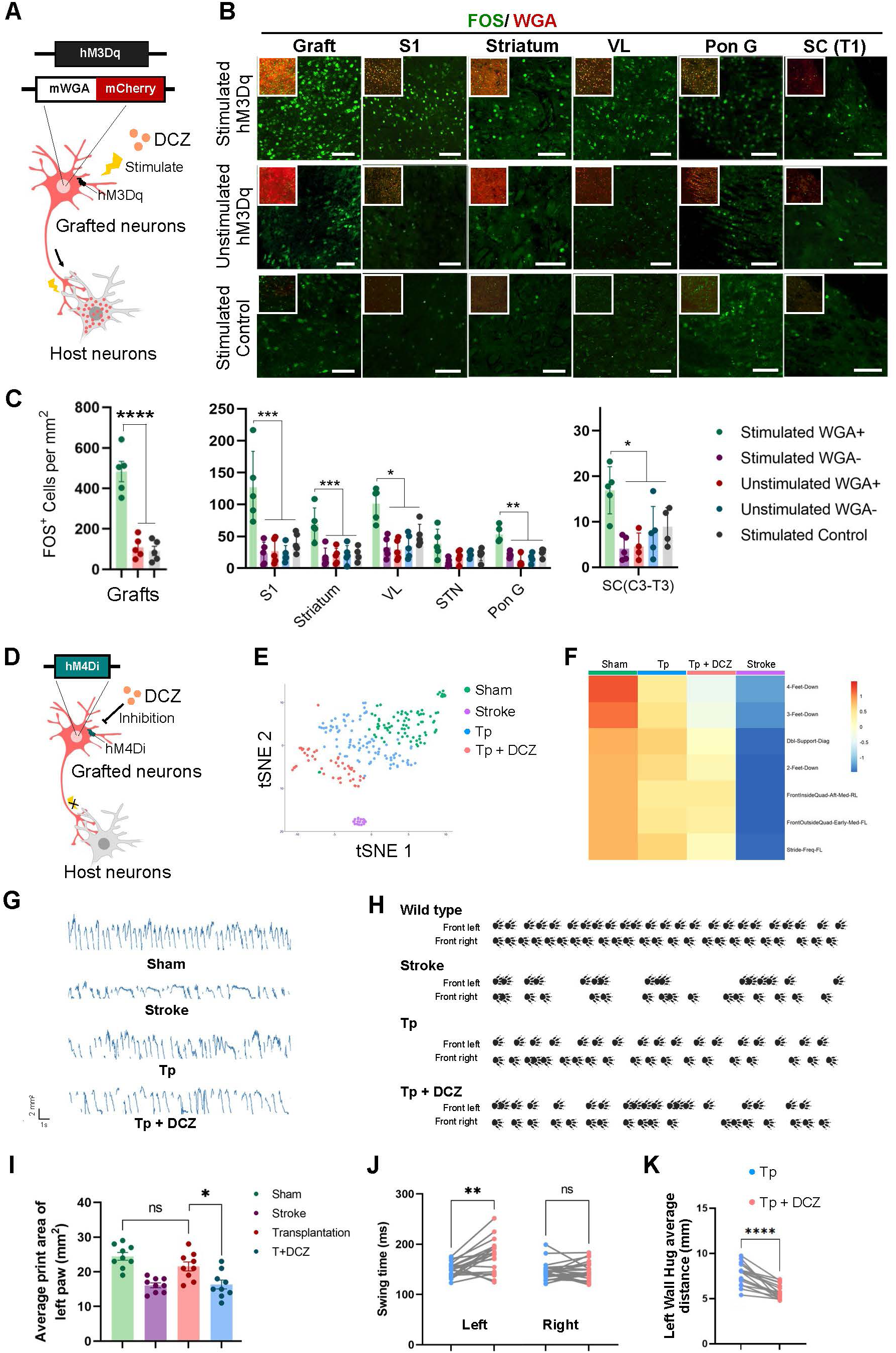
Transplanted cortical neurons make functional connections with host neurons. (A) Cartoon outlining the strategy to test the functional connections between the hESC-derived neurons and host neurons by activating the grafted DREADD/mWmC-human neurons with DCZ and examining the FOS expression in (WGA+) synaptically connected mouse neurons. (B) Immunostaining for FOS and WGA showing the expression of FOS in WGA+ neurons in the presence (top) or absence (middle) of DCZ at 12-month after transplantation. The bottom panels showing tissues from the animals grafted with hESC-derived neurons (without DREADD) following DCZ treatment (stimulated control). Scale bars, 100μm. (C) Quantification of FOS-expressing cells in the grafts (left), host brain (middle) and spinal cord (right) in stroke mice, stimulated or non-stimulated. *p<0.05, **p<0.01, ***p< 0.005, ****p<0.001, two-way ANOVA test. (D) Schematic showing the strategy to test the functional connections by inhibitory DREADD (hM4Di) hESC-derived neurons at 12-month after transplantation. (E) tSNE visualization of clustering of gait analysis for sham, stroke, transplanted (Tp), and hM4Di activated (Tp+DCZ) mice. (F) Heatmap showing difference in parameters among the sham, stroke, transplanted, and transplanted with hM4Di activation group from the stroke group. (G) Traces of the front left paws touching the treadmill belt in sham, stroke, Tp, and Tp+DCZ groups. (H) Footprints of the front paw captured in sham, stroke, Tp, and Tp+DCZ mice. (I) Quantification of average print area of the contralateral front paw in the four groups. n=9 mice in each condition. *p=0.0253, two-sided Student’s t-test. (J) Quantification of swing time of the left and right front paw before and after DCZ treatment in the transplanted mice. **p=0.0031, paired two-sided Student’s t-test. (K) Quantification of the average distance of left wall hug before and after hM4Di activation in the transplanted mice. ****p=0.0002, paired two-sided Student’s t-test. All Data are presented as mean±SEM.

To investigate whether the neural circuits regulated by transplanted neurons restore motor function in the stroke mice, we used NPCs from an inhibitory hM4Di DREADD hESC line as donors (Figure 3D and S8B-S8F). Gait analysis by TreadScan showed that cell transplantation improved motor function in stroke mice (Figures 3E and 3F). However, after DCZ administration to inhibit the transplanted neurons, the motor functions significantly declined, indicated by reduced time and frequency of the front left paws (contralateral side) touching the treadmill belt (Figures 3F and 3G). Footprint analysis of the front left paw (FL) revealed that inhibiting the transplanted neurons led to a recurrence of motor dysfunction in the previously recovered left limb (Figures 3H and 3I). Upon grafted neuron inhibition via hM4Di activation, there was a significant increase in the swing time of the left paw, while no notable changes were observed in the right paw (Figure 3J). Meanwhile, the mice stayed close to the contralateral wall after DCZ treatment (Figure 3K). To establish the specificity of the functional effects, we conducted a detailed TreadScan analysis of locomotor patterns across all four paws. We found that recovery was most pronounced in the front left paw (FL), and DCZ administration, which inhibits the transplanted neurons, resulted in a significant decrease in FL motor performance (Figure S9A). In contrast, the other three paws did not show DCZ-related changes (Figure S9B). These findings demonstrate that DCZ specifically impaired left forelimb function, consistent with our observation that transplanted cortical neurons projected to the spinal cord at the T3 level and formed synaptic connections with lower motor neurons (Figure S9C). These data suggest that the transplanted neurons form functional connections with the host neurons, and the neural circuits repaired by transplanted cortical neurons underline the motor function restoration.

### Projection of transplanted cortical neurons is subtype-specific in the ischemic brain

Transplanted cortical neurons extended axons to multiple targets, prompting a critical question of how the axonal targeting is regulated in the injured adult brain. To address this, we combined multiple retrograde tracing with snRNA sequencing (MRT-seq)^5^ to reveal the transcriptional profiles of grafted neurons projecting to specific regions. Four retrograde AAVs (GFP⁺), each carrying a unique barcode, were injected into the primary somatosensory cortex (S1), striatum, thalamus, and medulla (MY) five weeks prior to graft dissection for snRNA-seq (Figure 4A). The retro-AAV broadly labelled the L5 and L6 host cortical neurons (Figure S10A), as expected. The transplanted neurons, labelled by GFP (Figure 4B) and collected from three animals for snRNA-seq (Figures S10B and S10C). Strikingly, the retrogradely labelled grafted neurons formed distinct clusters (Figure 4C), indicating that projection-defined neuronal groups correspond to specific transcriptional subtypes. Cortical projection neurons are commonly classified into four subtypes: extratelencephalic (ET), intratelencephalic (IT), corticothalamic (CT), near-projecting (NP) neurons^22^, although snRNA-seq revealed at least 100 putative cortical neuronal subclasses^23^. Alignment with a public dataset of adult human M1 cortex (Figures S10D and S10E) revealed that retrogradely labelled cells partially correspond to four neuronal subtypes: L5 ET, L5 IT, L6 CT, and L5/6 NP (Figures 4C and 4D). Therefore, we named the grafted cortical neurons as grafted L5 intratelencephalic (G-L5 IT, striatum-projecting), grafted L6 CT (G-L6 CT), grafted L5 extratelencephalic (G-L5 ET, medulla-projecting), and grafted L5/6 near-projecting (G-L5/6 NP) neurons.

**Figure 4.**
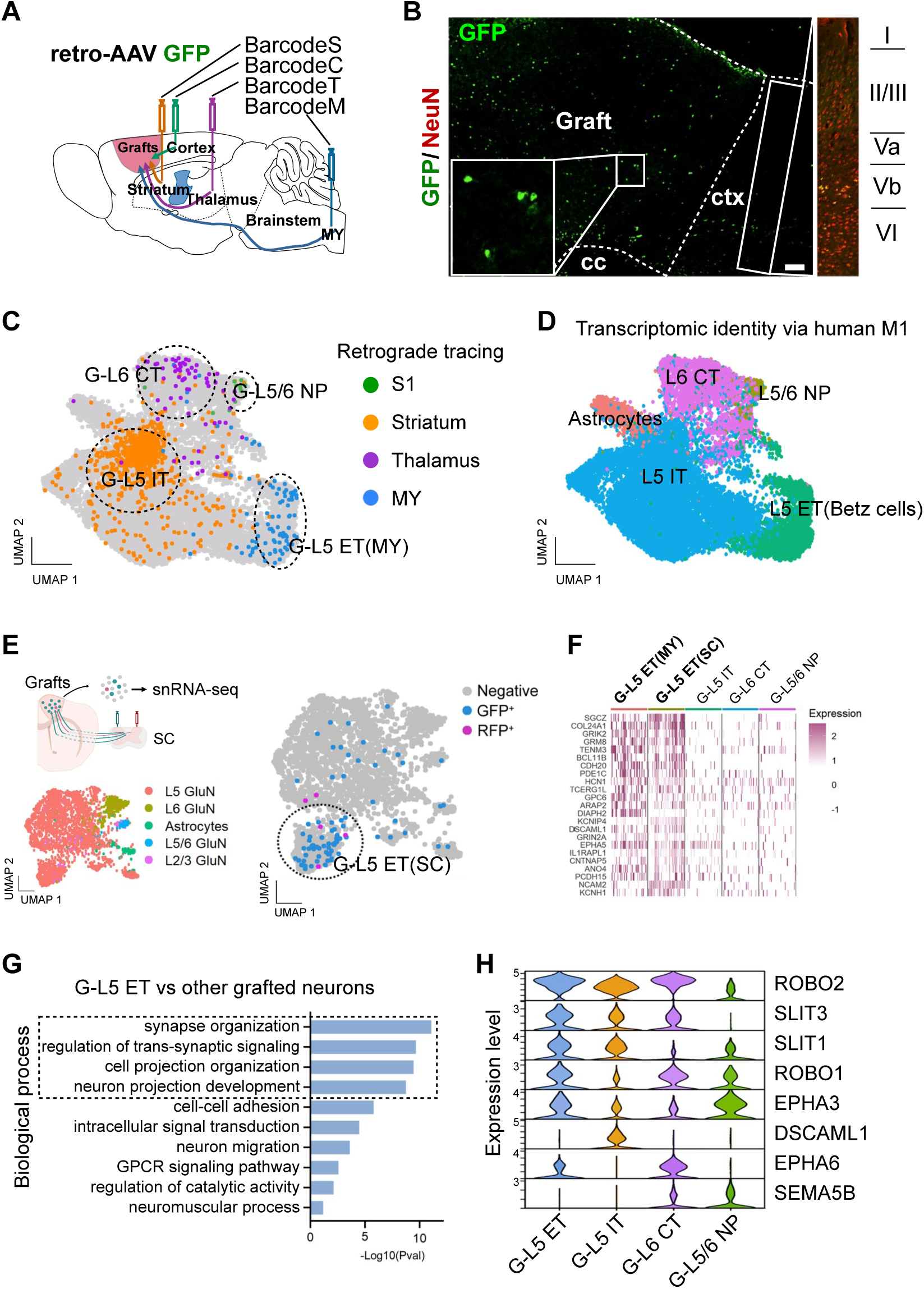
Transplanted cortical neuronal subtypes project to specific brain regions revealed by MRT-seq. (A) Cartoon showing the strategy for multiple barcoded retrograde tracing by injecting AAV into the somatosensory cortex, striatum, thalamus, and medulla of transplanted mice (left). (B) Immunostaining showing GFP labelling in the grafts and the neighboring host cortex. The inset is magnified on the right showing NeuN and GFP-labeled cells in a layered structure. Scale bars, 100μm. (C) UMAP visualization of clustering of retro-AAV labeled nuclei in the grafts in (A). (D) UMAP visualization showing the subtypes of transplanted cells using the human M1 snRNA-seq reference dataset. L5 ET, layer V extratelencephalic neurons; L5 IT, layer V intratelencephalic neurons; L6 CT, layer VI corticothalamic neurons; L5/6 NP; layer V/VI near-projecting neurons. (E) UMAP visualization of grafted neurons retrogradely labelled from the SC. The experimental strategy and UMAP visualization of transplanted neuronal subtypes are shown in the left panel. UMAP visualization of transplanted neurons labelled with retro-AAV is presented in the right panel. GluN, glutamatergic neurons; Astrocytes, astrocyte lineage cells; L2/3, layer II or III; L5, layer V; L6, layer VI; L5/6, layer V or VI. (F) Heatmap showing the top L5 ET marker expression of 50 randomly selected nuclei in each subtype of transplanted neurons. (G) Bar plot of enriched Gene Oncology (biology process) terms of G-L5 ET (MY) in the graft dataset (C). (H) Violin plot of canonical axon guidance genes for each subtype of transplanted cortical neurons in (C). S1, primary somatosensory cortex. MY, medulla. SC, spinal cord.

To identify the transcription profiles of G-L5 ET neurons projecting to the SC, we injected retrograde AAV with GFP in the contralateral SC and retro-AAV with RFP in the ipsilateral SC, both at the C7-T1 segments (Figures S10F-S10I). The host neurons labelled by AAV in the intact cortex were located in L5 (Figure S10H). 97% of retrogradely labelled grafted neurons projected to the contralateral SC (GFP^+^) whereas only 3% to the ipsilateral side (RFP^+^), as indicated by snRNA-seq (Figure 4E). Over 86% of all labelled cells belonged to the same neuronal cluster, and it shared a similar transcriptome with G-L5 ET (Figure 4F), including HCN1, KCNH1, BCL11B, SGCZ, GRIK2, and CNTN3. In particular, HCN1 channel-dependent membrane properties can distinguish L5 ET (human Betz cells) from L5 IT neurons^24^. Betz cells are characterized by low input resistance and distinct peak resonance, reflecting their large size and high expression of genes associated with the HCN channel^24,25^. The projection pattern of G-L5 ET neurons in the SC is also similar to that of human Betz cells (Figure S10J). To explain the relatively human-specific connections in rodents, we analyzed the DEGs of synaptic assembly between human and mouse corticospinal neurons. A specific combination of synaptic assembly genes, including ROBO2, NRCAM, and L1CAM, was significantly higher in the G-L5 ET neurons compared to mouse corticospinal neurons (Figure S10K). ROBO2, NRCAM, and L1CAM are critical receptors on the growth cone to connect to lower motor neurons^26–28^. They also play a vital role in commissural neurons in forming the decussation of the pyramids.

These data indicate substantial transcriptomic differences among the four projection defined subtypes of transplanted neurons. To investigate how transcriptomic differences influence the projection patterns of these neurons in the adult brain, we analyzed the DEGs among transplanted neurons with distinct projection targets. Gene ontology (GO) analysis of DEGs in G-L5 ET, G-L5 IT, G-L6 CT, and G-L5/6 NP neurons revealed that the top GO terms of biological processes were associated with axon guidance and synapse organization/transmission (Figures 4G and S10L-S10N). Among the four projection-defined subtypes, each expressed distinct combinations of canonical axon guidance genes, such as ROBO1, ROBO2, SLIT1, SLIT3, and EPHA3 (Figure 4H). These results highlight the transcriptomic predominance of specific genes related to axonal projection, explaining why transplanted neurons of one specific transcriptomic subtype mainly project to the same brain region.

### The projection pattern is regulated by transcription factors

We then ask whether modifying neuronal subtypes or their corresponding transcriptional codes can alter their projection pattern in the disease model. Ctip2 is an essential transcriptional factor in L5 cortical neurons^29,30^. We generated a CTIP2 knockdown (CTIP2-kd) hESC line and transplanted CTIP2-kd NPCs into the ischemic cortex (Figure 5A). Interestingly, the axons of CTIP2-kd neurons primarily projected to the amygdala (Amy) through the stria terminalis (ST) in the ipsilateral hemisphere (Figures 5B and 5C), with minimal projections to the subcortical regions (Figures 5B and S11A-S11C). Quantification of projections showed that the projection pattern of CTIP2-kd neurons was significantly different from that of wild-type neurons with more to hippocampus, basolateral amygdaloid nucleus (BLA), and basomedial amygdaloid nucleus (BMA) but less to subcortical regions (Figures 5D and 5E). Furthermore, robust axons from transplanted CTIP2-kd neurons crossed to the contralateral cortex through the corpus callosum (cc), which is rarely observed in wild-type ESC-derived neurons (Figure 5F). In germline Ctip2-deficiency mice, although the L5 neurons lack robust axons along the CST, they still project to subcortical regions and do not cross to the contralateral hemisphere^29^. The distinct projection patterns from grafted CTIP2-kd grafted neurons indicate the differences in axon guidance between the self-organizing and the mature brain environment.

**Figure 5.**
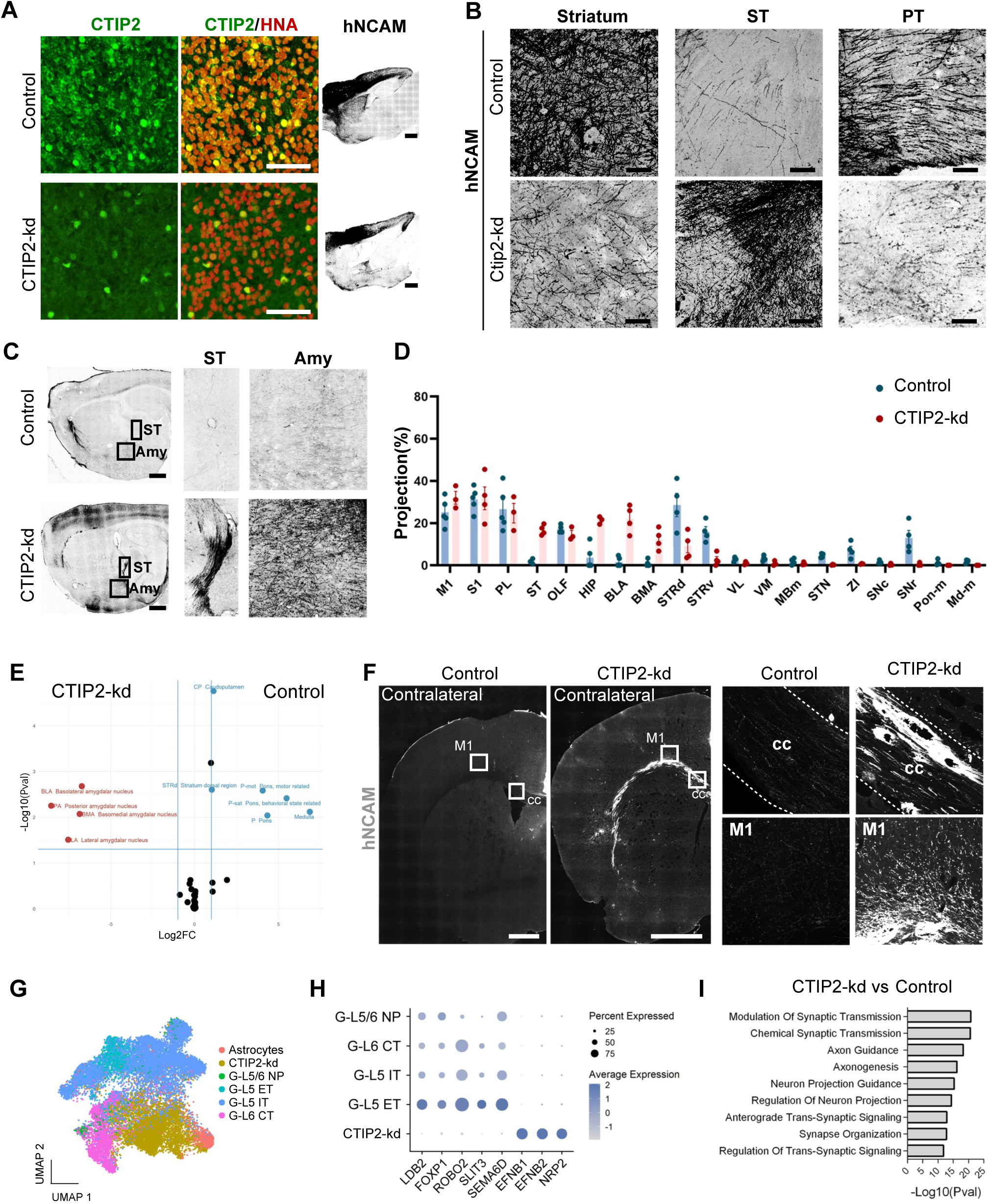
Grafted human cortical neurons alter the projection pattern when CTIP2 is knocked down. (A) Immunostaining for CTIP2 and HNA showing that reduction of CTIP2 expression in CTIP2-kd (bottom) than the wild-type (top) graft. Scale bars, 100μm. Right panels showing the hNCAM immunostaining on sagittal brain slices. Scale bars, 1mm. (B) Immunostaining for hNCAM showing graft-derived axons of control cell line (top) and Ctip2-kd cell line (bottom) in the striatum, ST, and PT in the ipsilateral hemisphere of stroke mice. (C) Immunostaining for hNCAM on the sagittal brain slices showing axon projection of the transplanted neurons derived from the control cell line (left) and CTIP2-kd cell line (right) in the stria terminalis (ST) and amygdala (Amy). (D) Quantification of axon projection of grafted neurons in ipsilateral brain regions. n=4-5 mice in each group. (E) Scatter plot showing brain regions with axon projection by the control cell and CTIP2-kd cell line. X: log2 fold change of axon projection, Y: –log10(p-value). (F) Immunostaining for hNCAM on the coronal sections showing axon projection of transplanted neurons derived from the control and CTIP2-kd cell line. Right panels showed the magnification in the cc, M1. Scale bars, 1mm. (G) UMAP visualization showing the subtypes of transplanted cells derived from control and CTIP2-kd cell lines after Seurat integration. (H) Dot plot of top marker genes of control and CTIP2-kd neurons showing the different expression levels of axon guidance genes. (I) Bar plot of enriched Gene Oncology – Biology Process terms between CTIP2-kd neurons versus control neurons. All Data are represented as mean±SEM

To reveal how CTIP2-kd alters axonal pathfinding, we integrated the snRNA profiling of control and CTIP2-kd cells and found that while their transcriptome distance is close, 76% of CTIP2-kd cells were not identified as any wild-type grafted neuron subtype (Figures 5G and S11D), indicating modified neuronal subtypes. The CTIP2-kd neurons also lack the markers of wild-type grafted neurons, such as LDB2 and FOXP1 (Figure 5H). The top-10 GO terms of biological process are linked to axon guidance and connectivity (Figure 5I). Among them, 61.5% of DEGs are related to axon projection (Figure S11E). In particular, the RNA levels of the axonal guidance molecules normally expressed in L5 ET neurons, such as ROBO2 and SLIT3, were downregulated whereas EFNB1, EFNB2, NRP2 were upregulated after knocking down CTIP2 (Figure 5H). ROBO2 and SLIT3 are required for forming the subcortical projection while EFNB1, EFNB2, and NRP2 play critical roles in guiding axons toward the amygdala and the contralateral hemisphere, which may explain the altered axonal trajectory.

Moreover, paw-resolved grid-walking and TreadScan analysis revealed that knockdown of a single transcription factor CTIP2 in transplanted cortical neurons led to strikingly divergent recovery profiles (Figure S11F and S11G). Mice receiving normal CTIP2^+^/FEZF2^+^ neurons showed recovery predominantly in the contralateral forelimb (Figure S11F and S11H), whereas mice transplanted with CTIP2-KD neurons showed recovery largely confined to the ipsilateral hindlimb (Figure S11F-S11I). Notably, these behavioral outcomes are consistent with the anatomical projections of the transplanted neurons within the CNS, i.e., the CTIP2-kd neurons projected to the contralateral cortex to restore the corresponding functions. Hence, modification of the transcriptome by altering a single transcription factor such as CTIP2 changes the axonal trajectory, highlighting the requirement of a special code for the same neuronal type to navigate in the mature brain.

### Grafted neuronal subtypes carry unique transcriptional codes for axonal projection

The above analysis revealed the match between the transplanted cortical neuronal subtypes and the expected projection targets, raising the intriguing question of how hPSC-derived neurons navigate and establish connectivity within the disorganized environment of the injured brain, in contrast to the self-organizing processes during development. We first evaluated the maturation state of the grafted neurons by comparing them with human cortical neurons at different stages (Figure 6A)^24,31^. The clusters of grafted neurons predominantly overlap with adult deep-layer cortical neuronal types (Figure S12A), while fewer clusters overlap with postnatal neurons (Figure S12B). The similarity was further validated by pseudotime analysis (Figures S12C-S12F). Our data indicate that the four subtypes of 12-month grafted human neurons transcriptomically resemble the corresponding subtypes of adult human cortical neurons in the M1 (Figure 3A). Both G-L5 ET and H-L5 ET neurons possess significantly higher expression levels of ROBO2, PTPRM, and SEMA3D (Figures 6A and S12G), which likely facilitate the formation and maintenance of corticospinal connections with spinal motor neurons^24,32^. Interestingly, several axon guidance molecules, including DRAXIN, UNC5C, FLRT2, and NEO1, are more highly expressed in grafted corticospinal neurons compared to endogenous human neurons (Figure S12G). These molecules are associated with axonal repulsion, suggesting that stronger repulsion-mediated pruning may be essential for directing axons along correct trajectories in the adult brain, which is also observed in L5 IT, L6 CT, and L5/6 NP neurons (Figure 6A). This may help explain why transplanted neurons, although closely mimicking endogenous neurons in function, often fail to extend their axons to appropriate target regions.

**Figure 6.**
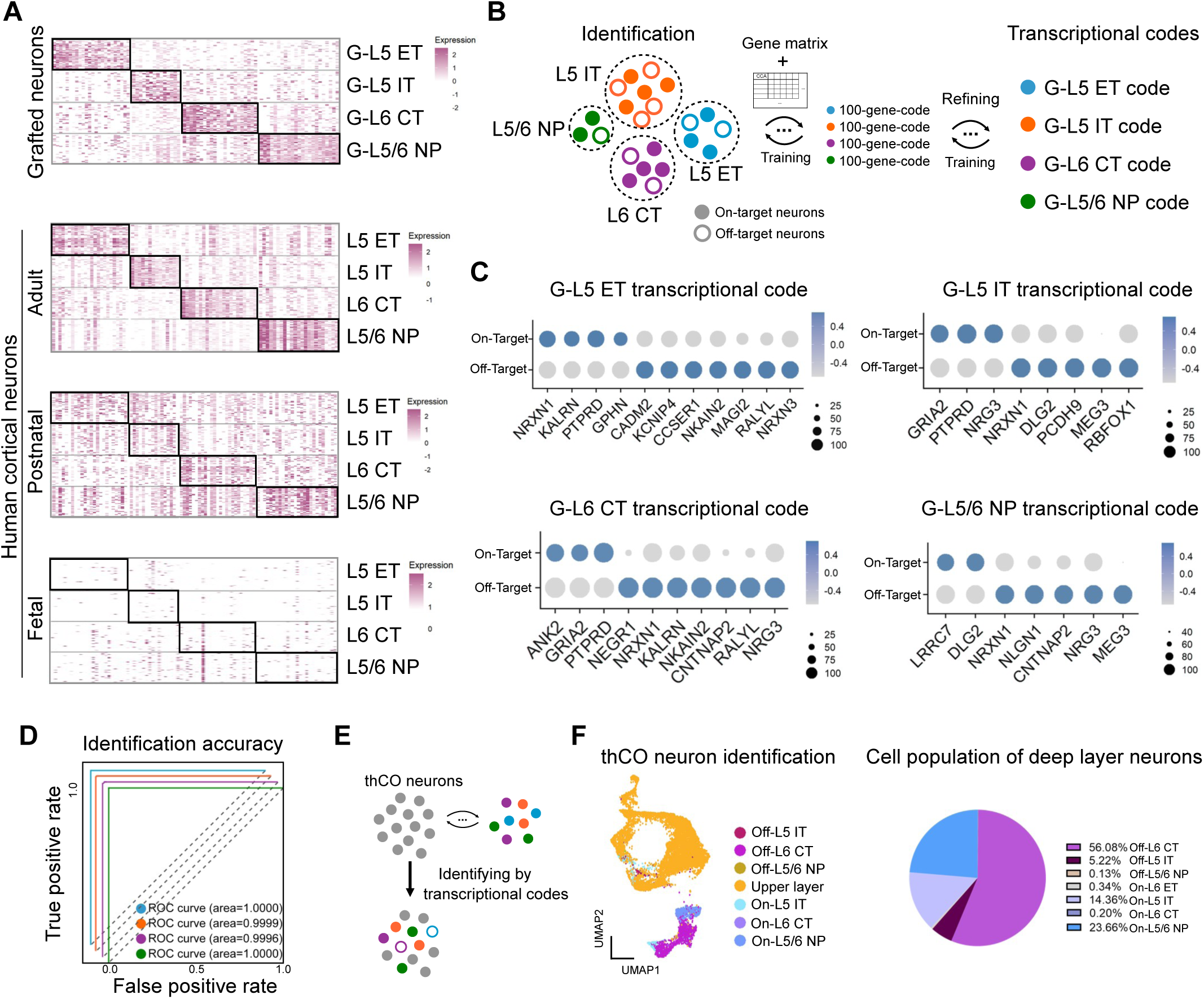
Transcriptional codes for grafted cortical neurons. (A) Heatmap showing the top marker expression in 50 randomly selected nuclei in each subtype of transplanted, adult, postnatal (8-month), fetal (6-month) human cortical neurons. (B) Schematic showing the identification of transcriptional codes with machine learning for grafted neurons. (C) Dot plot showing the expression levels of the minimal number of genes (transcriptional codes) to identify on-target grafted neurons in four subtypes. (D) ROC curves showing the identification accuracy of grafted neurons in each subtype. (E) Strategy for identification of cortical neurons from transplanted human cortical organoid (thCO). (F) UMAP visualization (left panel) and quantification (right panel) of publicly available data of transplanted cortical neurons identified by transcriptional codes. On-, On-target grafted neurons. Off-, Off-target grafted neurons.

We next sought to identify the transcriptional code for each of the subtypes that establish projections to the intended target regions (on-target), as opposed to those that merely exhibit transcriptomic profiles resembling their cortical counterparts without appropriate targeting (off-target), by comparing the transcriptomic differences between projection-defined subtypes and their human counterparts. A set of 100 genes was calculated from all genes using the “SelectKBest” method with the “f_classif” statistical test to define grafted neurons in each subtype and further refined using L1-regularized logistic regression and cross-validated by Bayesian Additive regression trees to obtain a code with minimal numbers of genes (Figure 6B). Although the gene expression profiles of one specific neuronal subtype are largely overlapping, our results demonstrate that differential expressions of several selected genes is sufficient as markers to distinguish on-target neurons from off-target neurons, serving as a form of transcriptional code for grafted neurons (Figures 6C and 6D). Across the four neuronal subtypes, these gene combinations exhibit distinct variations, suggesting the subtype specificity. Notably, most of these genes are implicated in axon guidance and synapse organization, including GPHN, KALRN, PTPRD, DSCAM, NRXN1, NRXN3, GRIA2, ANK2, NEGR1, NRG3, LRRTM4, LRRC7, MEG3, CADM2, DLG2, and CNTNAP2 (Figure 6C).

To validate our formulation, we analyzed the neuronal subtypes in the cortical organoids transplanted into the mouse S1 cortex^13^ by integrating the snRNA-seq reference dataset (Figure 6E). According to our transcriptional code-based identification, the neurons in the transplanted cortical organoid comprised of 0.34% G-L5 ET, 0.20% G-L6 CT, 23.66% G-L5/6 NP, and 14.36% G-L5 IT on-target neurons (Figure 6F). Indeed, the transplanted (organoid) neurons project predominantly to the ipsilateral cortical and striatal regions but rarely to the pyramidal tract and the ventrolateral/ventromedial thalamic nucleus^13^, supporting the accuracy of our model. Similarly, our codes reveal the lack of IT, ET, CT, and NP subtypes in the CTIP2-kd neurons (Figure S12H), consistent with the lack of ipsilateral hemisphere projections observed anatomically. Our findings show the critical role of specific axon guidance profiles in identifying the neuronal subtypes. These transcriptomes, while similar to those of the corresponding adult human neuronal subtypes, exhibit unique features in combination and/or expression levels, serving as transcriptional codes for the grafted neurons to find their targets in the adult CNS.

## DISCUSSION

Defining if and how the transplanted neurons find and make functional connections with their targets in the adult brain will enable us to develop effective strategies for and predict the outcome of cell therapy. We established the correspondences between projection targets and transplanted neuronal subtypes and identified the transcriptional code for specific axonal guidance and synapse assembly in the mature brain with machine learning-based regression analysis. These transcriptional codes associate transplanted neuronal identity with their projection patterns. This is further supported by findings that changes in neuronal identity induced by transcription factors such as CTIP2 result in profound alterations in axonal projections and target recognition, suggesting that the projection directions of transplanted neurons in the mature brain can be modulated through transcription factor regulation. The circuits reconnected by the transplanted human cortical neurons are specific and functional, as indicated by the mono-synaptic anterograde tracing and activity-dependent FOS expression in the target neurons, corresponding to motor behavioural recovery of the stroke mice. These transcriptome profiles serve as tags to identify the complex guidance combinations, setting the foundation for developing specific cell types for circuit repair.

It has been demonstrated that transplanted neurons form functional connections with host neurons, contributing to the therapeutic effects in animal models of neurological diseases^13,33–36^. The premise for choosing specific types of neuronal cells for transplantation is based on the information we have learned in developing animal models, i.e., a developing neuron finding its targets is largely determined by its specific transcriptiome^6,37–39^. However, the environment of the brain undergoes substantial modification after maturation, which alters the organization patterns of projections and connectivity. Hence, the axonal projection of grafted neurons to their long-distance targets is usually insufficient^2,3^, or establishes inappropriate connections^40^. It is thus desirable to identify the intrinsic properties or transcriptional codes of the transplantable neuronal cells to achieve high-fidelity circuit integration in the adult brain. Hence, we identified the specific combinations of axonal guidance and synapse organization gene profiles or codes for each cortical neuronal subtype. Notably, transplanted neurons exhibited a pronounced upregulation of repulsive guidance cues compared to endogenous neurons^24^. This differential expression pattern suggests the special balance between attractive and repulsive guidance signals required for axonal navigation in the adult brain. The drastic change in projection patterns by the grafted cells with CTIP2-kd highlights the regulatable properties of the grafted neurons in pathfinding and circuit integration even in the adult brain. And that circuit rewiring shifts functional contribution precisely from the ipsilateral forelimb to contralateral hindlamb. Indeed, the corticospinal neuronal subtype in our transplanted cells project axons via the pyramidal tract, cross over to the contralateral side at the medulla level, and reconnect with the spinal motor neurons, which corresponds to the motor behavioral recovery of the stroke mice. A striking feature of such connections is direct synaptic connection between the human cortical motor neurons and the mouse spinal motor neurons, a property in primates, besides indirect synapses through interneurons, a characteristic in rodents. Such a phenomenon again highlights the critical role of the specific transcriptional code of the grafted neurons in circuit integration. Thus, the transcriptome of the transplanted cortical neuron subtypes may serve as identity tags to predict their projections and connectivity, providing a guideline for selecting appropriate neuronal cell types for targeted circuit reconstruction.

### Limitations of the study

The high-fidelity circuit reconstruction through transplanted neurons is influenced by numerous factors. Our study revealed the roles of transcriptional codes of the major cortical neuronal subtypes in pathfinding and synaptic assembly in the mature brain. Analysis of additional neuronal types in other model systems will help determine if it is a general principle. The similarity to as well as the difference from their endogenous counterparts suggests the potential impact of the environmental signals in modulating the transcription of the grafted cells, which warrants investigation.

## RESOURCE AVAILABILITY

Microscopy data reported in this paper will be shared by the lead contact upon request.

Any additional information required to reanalyze the data reported in this paper is available from the lead contact upon request.

### Lead contact

Requests for further information and resources should be directed to and will be fulfilled by the lead contact, Su-Chun Zhang.

### Materials availability

Neural transplantation cocktail generated in this study will be made available on request, but we may require a payment and/or a completed materials transfer agreement if there is potential for commercial application.

### Data and code availability

The snRNA-seq datasets and machine learning code are available at the Dryad with accession number 4j0zpc8qd (Dataset DOI: 10.5061/dryad.4j0zpc8qd). Allen Human Motor Cortex Atlas is from the Allen Brain Atlas Data Portal (http://help.brain-map.org/display/devmouse/API). Any additional information required to reanalyze the data reported in this paper is available from the lead contact upon request.

## ACKNOWLEDGMENTS

These studies were supported in part by a Duke/Duke-NUS Pilot grant, National Medical Research Council of Singapore (MOH-000207, MOH-000212), Open Fund-Young Individual Research Grant (OFYIRG23jan-0031), and Khoo Postdoctoral Fellowship Award (Duke-NUS-KPFA/2021/0047). We thank Qiang Yuan, Ye Sing Tan, Xianfeng Tian, Chung Hock Lee, and Mynn Michele Dy Varela for dedicated technical assistance.

## AUTHOR CONTRIBUTIONS

Conception and design of work, SC.Z., and ZF.W.; stroke model, ZF.W., DY.Z., and F. Y; cell transplantation, ZF.W., DY.Z., and F. Y; cell culture, ZF.W., DY.Z., and F. Y; animal breeding, ZF.W. and DY.Z.; lentivirus production, ZF.W.; cell line generation: ZF.W. and P.J.K.; electrophysiology, SM.C.; data collection, ZF.W. and DY.Z.; data analysis and interpretation, ZF.W., DY.Z., and SC.Z.; drafting the article, ZF.W. and SC.Z..

## SUPPLEMENTAL INFORMATION

**Document S1. Figures S1–S12 and Table S1.**

**Video S1-S5. TreadScan tests, related to Figure 3, S9**

## LEGENDS

**Figure S1.** Characterization of human cortical neurons before and after transplantation. (A) Immunostaining shows co-expression of CTIP2 and FEZF2 in human cortical neurons prior to transplantation. Scale bars, 200μm. (B) Immunostaining for HNA, CTIP2, and TBR1 shows expression of deep layer marks in human cortical NPCs before transplantation. Scale bars, 200μm. (C) Immunostaining of DAPI and GFAP showing host astrocytes in the glial scar surrounding lesion core. Scale bars, 1mm. (D) Immunostaining of GFP and NeuN showing survival of GPF+ NPCs in the lesion at 14-day post-transplantation on the right panel. Scale bars, 1mm. (E) Whole-mount fluorescence image shows that the graft (mCherry+) filled the ischemic motor cortex at 12 months post transplantation. Scale bars, 1mm. (F) UMAP visualization of clustering of nuclei from grafted cells. (G) Feature plot showing glutamatergic neuron markers SYT1+ (left) and SATB2+ (right) nuclei from (F). (H) Feature plot showing deep layer neuron markers CTIP2+/FEZF2+ (left) and TBR1+ (right) cells. (I) Feature plot showing upper layer neuron marker POU3F2+ cells. (J) Feature plot showing GABAergic neuron marker GAD1+ cells. (K) Feature plot showing astrocyte markers AQP4+ (left) and SLC1A3+ (right). (L) Immunostaining for HNA, SOX9, and Iba1 on the brain slices showing astrocytes and microglia in the grafts at 12 months post transplantation. Scale bars, 200μm. (M) Feature plot showing oligodendrocyte marker OLIG1 cells. (N) Feature plot showing proliferating cell marker KI67.

**Figure S2.** Cell fates of transplanted neurons before and after transplantation. (A) UMAP showing the grafted cells before transplantation, 3 months post-transplantation, 9 months post-transplantation, and 12 months post-transplantation after Seurat integration. (B) Feature plots showing the expression of SOX2, NES, RBFOX3, and SATB2 of cells in (A). (C) UMAP showing the co-expression of SOX2, NES, RBFOX3, and SATB2 of cells in (A). (D) Barplots showing the proportion of cells expressing SOX2, NES, RBFOX3, and STAB2 in (A). (E) UMAP showing the expression of CASP9 and MKI67 of cells in (A). Bar plots on the right panel showing temporal changes in the expression ratios of CASP9 and MKI67 following transplantation. The ratios remained largely unchanged from 3 months post-transplantation onward.

**Figure S3.** Myelination and vascularization in the graft. (A) Immunostaining of hNCAM and oligodendrocyte marker MBP showing graft-derived axons are surrounded by mouse oligodendrocytes. The cross-sectional view is shown in panel (A’), and side view in panel (A’’). Yellow arrowheads indicate the relationship between MBP and hNCAM signals. Scale bars, 100μm. (B) Immunostaining with MBP (oligodendrocyte marker) and HNA showing no expression of MBP in transplanted cells. Scale bars, 100μm. (C) Immunostaining with STEM121 and endothelial marker Laminin on the brain slices of 12-month after transplantation, showing the mouse blood vessel endothelium grew into the graft. Scale bars, 500μm. 200μm in magnification.

**Figure S4.** Transplanted human cortical neurons rebuild the motor-related neural circuits. (A) Schematic diagram showing the planes for brain sections. (B to E) Immunostaining of hNCAM showing axon projections by the grafted neurons in Pons and MY (B), thalamus, PT and Pons (C), striatum, ic and cp (D), and S1 and striatum (E), in stroke mice 12 months post transplantation. Scale bars, 1mm. 100μm in magnification. (F) Immunostaining for hNCAM showing axon projections by the grafted neurons in the contralateral and ipsilateral hemispheres at 12 months after transplantation. Scale bars, 1mm. 100μm in the magnification. (G) Immunostaining for hNCAM showing graft-derived axons in the contralateral spinal cord C7 (top) and T3 (bottom) at 12 months after transplantation. Scale bars, 500μm. (H) Quantification of axon projection of grafted neurons in brain regions. All Data are presented as mean±SEM.

**Figure S5.** Validation of anterograde mono-synaptic tracer mWmC. (A) Schematic of generating GFP/mWmC cell lines to validate mWmC (left). Immunostaining for NeuN, mCherry, and GFP showing that mCherry(mWmC) is only detected in NeuN+ cells (right). Yellow arrowheads indicate primary neuron (GFP+/mCherry+); White arrowheads indicate NeuN+ cells; open arrowheads indicate NeuN-cells. Scale bars, 100μm. (B) Immunostaining of NF-L, PSD95, GFP, and mCherry on neural progenitors showing that the mWmC is not presented in GFP-/PSD95-neural progenitors. Dashed lines indicate GFP-/PSD95-cells. Scale bars, 20μm. (C) Schematic diagram showing that primary neurons (GFP+/mWmC+) project axons distally to downstream neurons (hM3Dq-mCherry), validating that mWmC works as monosynaptic tracer. Immunostaining of NF-L, PSD95, GFP, and mCherry on downstream neurons showing no puncta red signals are detected in the second-order neurons. Arrows indicate puncta red signals in the cytoplasm. (C’ and C’’) indicate the first neurons which attach GFP+ neurites from primary neurons; C’’’ indicates the second-order neurons. Scale bars, 20μm.

**Figure S6.** Anterograde trans-synaptic tracer mWmC *in vivo*. (A) Strategy for injection of mWmC-AAV in M1c of the mouse brain. Immunostaining showing mCherry (mWmC) detection in the S1, striatum, VL, Pon G and T1 (spinal cord) in the mouse CNS. Scale bars, 200μm. (B) Immunostaining showing mCherry (mWmC) detection in the T1, T10, and L6 (spinal cord). Scale bars, 100μm.

**Figure S7.** Transplanted neurons synapse with specific target neurons. (A) Immunostaining for WGA and Ctip2 showing that host medium spiny neurons (Ctip2+) were labelled by WGA in the striatum at 12-mpt. Right panels show magnified view of the insets in the dorsal (A’) and ventral (A’’) striatum. Yellow arrowheads indicate WGA+/Ctip2+ neurons. Scale bars, 100μm. (B) Immunostaining with NeuN, WGA, and GAD67 showing that GABAergic neurons in the cortex and thalamus were WGA positive. Right panels show magnification of VL and S1. Yellow arrowheads indicate WGA+/GAD67+/NeuN+ cells. Scale bars, 1mm. (C) Immunostaining for vGLUT1 and WGA showed that WGA labels (vGLUT1+) glutamatergic neurons in the VL and STN. The magnification was shown in the right panels. Yellow arrowheads indicate WGA+/vGluT1+ cells. White arrowheads indicate WGA+/vGluT1-cells. Scale bars, 100μm. (D) Immunostaining showing no overlap of WGA with hostdopaminergic neurons (TH+) in the substantia nigra. White arrowheads indicate WGA+/ TH-cells. White arrows indicate TH+/WGA-cells. Scale bars, 100μm. (E) Immunostaining showing GAD67-expressing WGA-labeled host neurons in the spinal cord. The insets are magnified on the right, yellow arrowheads indicate WGA+/ GAD67+ cells. Scale bars, 100μm. (F) Quantification of the ratio of host neurons receiving output connections of grafted cells in different brain regions. *p<0.05, **p<0.01, two-sided Student’s t-test. Data are presented as mean±SEM.

**Figure S8.** Functional tests of DREADD/mWmC ESC-derived neurons. (A) Strategy for generation of hM3Dq-mCherry–mWmC-eGFP cell line (left). Immunostaining showing expression of GFP and mCherry(mWmC) in 2D-cultured neurons 60 (right top) and 90 (right bottom) days of induction. Scale bars, 100μm. (B) The left panel shows the schematic of hM4Di (DREADD)/mWmC neurons following DCZ treatment (inhibition). The middle panel shows the experimental strategy. The right panel shows the calcium imaging of upstream neurons and their downstream neurons following DCZ treatment. Scale bars, 100μm. (C) Heatmap showing the neuronal activity of mWmC^+^ and mWmC^-^ neurons after DCZ addition. The top panel showing the upstream neurons. The bottom panel showing the downstream neurons. The right panel showing the average intensity over time. (D) Heatmap showing neuronal activity of mWmC+ neurons in the striatum of mice transplanted with DREADD neurons after DCZ/PBS administration. The right panel showing the average intensity over time. (E) Immunostaining for FOS showing the expression of FOS in host neurons in the presence of DCZ at 12-month after hM4Di neural transplantation. Scale bars, 200μm. (F) Quantification of FOS-expressing cells in the host brain and spinal cord in stroke mice. *p<0.05, **p<0.01, Student’s t-test.

**Figure S9.** Association between limb movement and spinal motor neuron connections. (A and B) Heatmap of parameters of front left paw (FL), front right paw (FR), rear left paw (RL), and rear right paw (RR) in treadscan assessment of mice from transplantation (Tp), transplantation with DCZ injection (Tp + DCZ; inhabitation), and Stroke group. (A) shows FL. (B) shows FR, RL, and RR. (C) Immunostaining for mCherry and ChAT showing the synaptic connections between grafted neurons and spinal motor neurons in C7 and T2. Scale bars, 1mm.

**Figure S10.** Retrograde tracing for transplanted neurons. (A) Strategy for injection of retrograde-AAV GFP with barcodes in the transplanted mouse brain (left). Immunostaining for GFP and Ctip2 showing retrograde-AAV labelled layer V neurons in the intact cortex (right). Scale bars, 200μm. (B) Strategy for MRT-seq at 5 weeks after AAV injection. UMAP showing clusters of nuclei from transplanted cells extracted from the mouse brain with barcoded retro-AAV-GFP injection. (C) Heatmap showing marker expression of clusters in (B). (D) UMAP showing the grafted cells and human M1c snRNA-seq reference dataset after Seurat integration. (E) UMAP shows clustering of nuclei from grafted cells using label transferred from the human M1c snRNA-seq reference dataset. (F) Strategy for retrograde AAV injection in the spinal cord of transplanted stroke mice at 12 months after transplantation. (G) Immunostaining for RFP and NeuN showing retrograde AAV-RFP-labelled axons in the ipsilateral spinal cord of transplanted stroke mice. Scale bars, 1mm. (H) Immunostaining of retrograde-AAV-GFP and RFP-labeled neurons in the cortex of stroke mice. Scale bars, 200μm. (I) Immunostaining for GFP and HNA showing the transplanted neurons were labelled by retro-AAV. Scale bars, 200μm. (J) Immunostaining showing axonal projection (hNCAM+) and output connection (mCherry+) of grafted cells to the spinal cord (C8). DH, dorsal horn. D, dorsal funiculus. VH, ventral horn. L, lateral funiculus. Scale bars, 500μm. (K) Dot plot of top axon guidance/synapse assembly genes of human L5 ET versus mouse L5 ET neurons. The genes expression on Grafted L5 ET neurons is represented at the bottom of the plot for top feature genes. (L-N) Bar plot of enriched Gene Oncology – Biology Process terms of cluster G-L5 IT (L), G-L6 CT (M), and G-L5/6 NP (N) in the graft dataset.

**Figure S11.** The projection pattern of transplanted CTIP2-kd neurons. (A) Schematic for brain sections of mice transplanted with CTIP2-kd human neurons. (B-C) Immunostaining of hNCAM showing axon projections by the grafted CTIP2-kd neurons in the striatum and PT (B), SI, ic and cp (C) at 12 months post transplantation. Scale bars, 1mm. (D) UMAP visualization shows the subtypes of transplanted cells derived from control and CTIP2-kd cell lines after Seurat integration. (E) Network of enriched Gene Oncology – Biology Process terms of cluster CTIP2-kd neurons versus control neurons. (F) Barplot showing the proportion of paw faults over number of steps in grid walk assessment of mice with CTIP2-kd/hM4Di neurons transplantation, CTIP2-kd/hM4Di neurons transplantation with DCZ injection and with WT neurons transplantation. (G) UMAP showing clustering of treadscan assessments of mice from 6 groups: Sham, Stroke, Tp (transplantation), Tp + DCZ (WT neurons transplantation with DCZ injection), CTIP2-kd (CTIP2-kd/hM4Di neurons transplantation), CTIP2-kd + DCZ (CTIP2-kd/hM4Di neurons transplantation with DCZ injection). (H and I) Heatmap of parameters of front left paw (FL), front right paw (FR), rear left paw (RL), and rear right paw (RR) in treadscan assessment of mice from Tp, Tp + DCZ, CTIP2-kd, CTIP2-kd + DCZ and Stroke group. (H) shows FL. (I) shows FR, RL, and RR.

**Figure S12.** Comparison of transplanted neurons with endogenous human neurons. (A and B) Heat maps of grafted cell clusters overlap by RNA-seq integration with human adult (A) and postnatal (B) cortical cell clusters. (C) UMAP plots showing the maturation trajectory in each subtype of human cortical neurons from fetal, postnatal, and adult cortex. (D) UMAP plots showing the transplanted, postnatal, and adult cortical neurons after Seurat integration. (E) Quantification of pseudotime in each subtype of cortical neurons. (F) Bar plot showing the difference in pseudotime distance between grafted neurons and postnatal versus adult neurons, indicating their relative similarity. (G) Dot plot of canonical guidance gene expression on G-L5 ET neurons. The genes expression on human L5 ET neurons (H-L5 ET) is represented on the top of the plot for top feature genes. (H) Dot plot showing the profiles of the minimal number of genes required to identify CTIP2-kd neurons from on-target grafted neurons in four subtypes.

