## Supplemental Figures for "Coding for Circuit Integration in the Injured Brain by Transplanted Human Neurons"

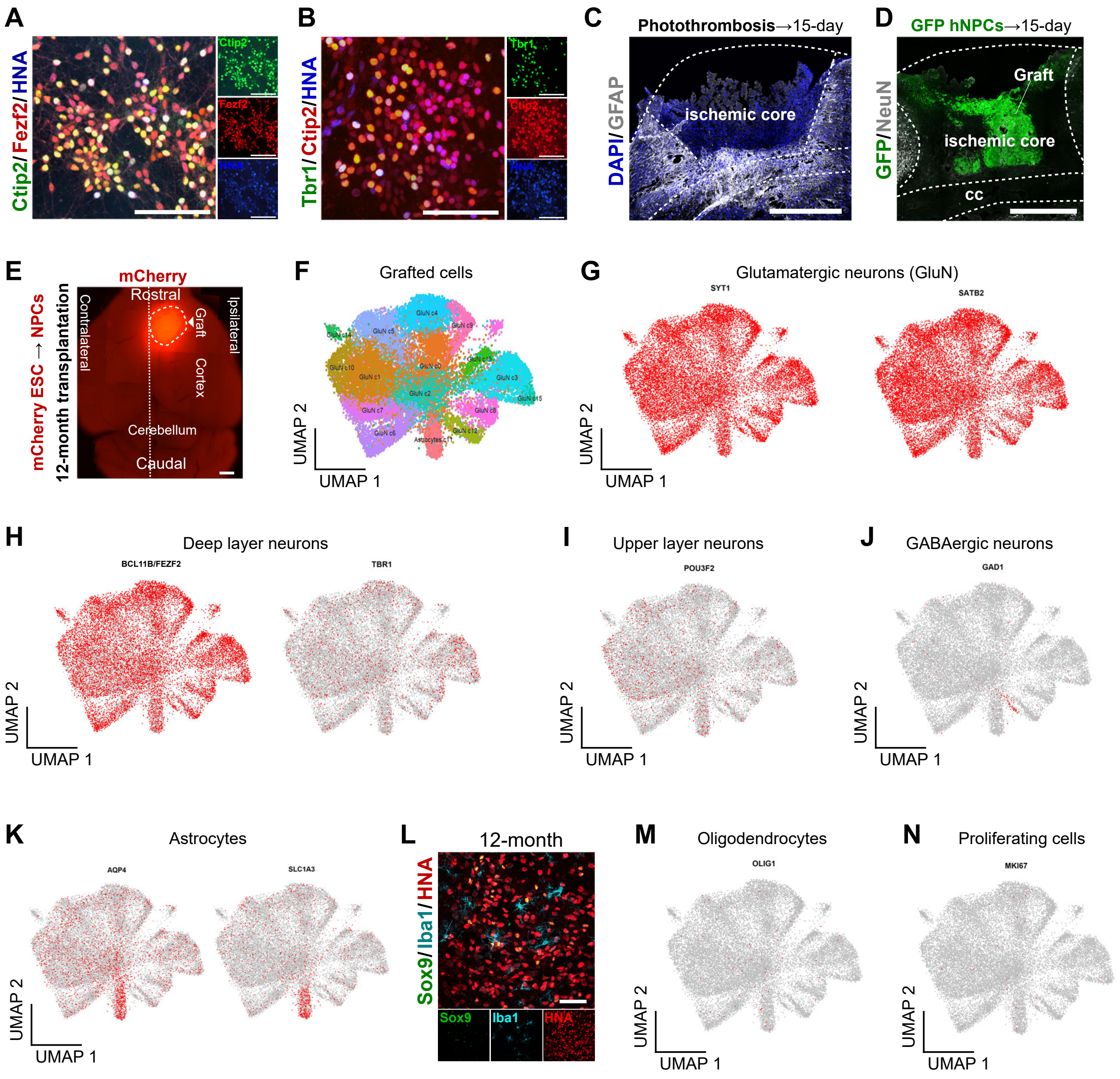


**Figure S1. Characterization of human cortical neurons before and after transplantation.**

(A) Immunostaining shows co-expression of CTIP2 and FEZF2 in human cortical neurons prior to transplantation. Scale bars, 200μm.

(B) Immunostaining for HNA, CTIP2, and TBR1 shows expression of deep layer marks in human cortical NPCs before transplantation. Scale bars, 200μm.

(C) Immunostaining of DAPI and GFAP showing host astrocytes in the glial scar surrounding lesion core. Scale bars, 1mm.

(D) Immunostaining of GFP and NeuN showing survival of GPF+ NPCs in the lesion at 14-day post-transplantation on the right panel. Scale bars, 1mm.

(E) Whole-mount fluorescence image shows that the graft (mCherry+) filled the ischemic motor cortex at 12 months post transplantation. Scale bars, 1mm.

(F) UMAP visualization of clustering of nuclei from grafted cells.

(G) Feature plot showing glutamatergic neuron markers SYT1+ (left) and SATB2+ (right) nuclei from (F).
(H) Feature plot showing deep layer neuron markers CTIP2+/FEZF2+ (left) and TBR1+ (right) cells.
(I) Feature plot showing upper layer neuron marker POU3F2+ cells.
(J) Feature plot showing GABAergic neuron marker GAD1+ cells.
(K) Feature plot showing astrocyte markers AQP4+ (left) and SLC1A3+ (right).
(L) Immunostaining for HNA, SOX9, and Iba1 on the brain slices showing astrocytes and microglia in the grafts at 12 months post transplantation. Scale bars, 200μm.
(M) Feature plot showing oligodendrocyte marker OLIG1 cells.
(N) Feature plot showing proliferating cell marker KI67.

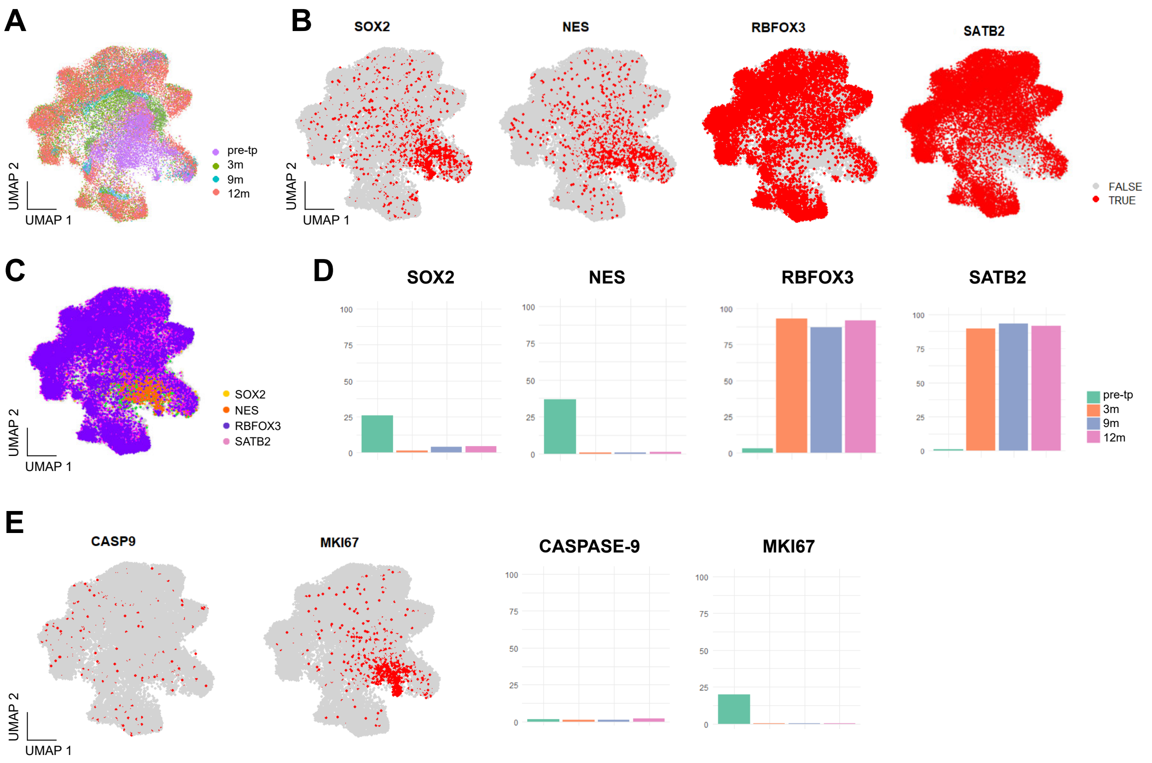


**Figure S2. Cell fates of transplanted neurons before and after transplantation.**

(A) UMAP showing the grafted cells before transplantation, 3 months post-transplantation, 9 months post-transplantation, and 12 months post-transplantation after Seurat integration.

(B) Feature plots showing the expression of SOX2, NES, RBFOX3, and SATB2 of cells in (A).

(C) UMAP showing the co-expression of SOX2, NES, RBFOX3, and SATB2 of cells in (A).

(D) Barplots showing the proportion of cells expressing SOX2, NES, RBFOX3, and STAB2 in (A).

(E) UMAP showing the expression of CASP9 and MKI67 of cells in (A). Bar plots on the right panel showing temporal changes in the expression ratios of CASP9 and MKI67 following transplantation. The ratios remained largely unchanged from 3 months post-transplantation onward.

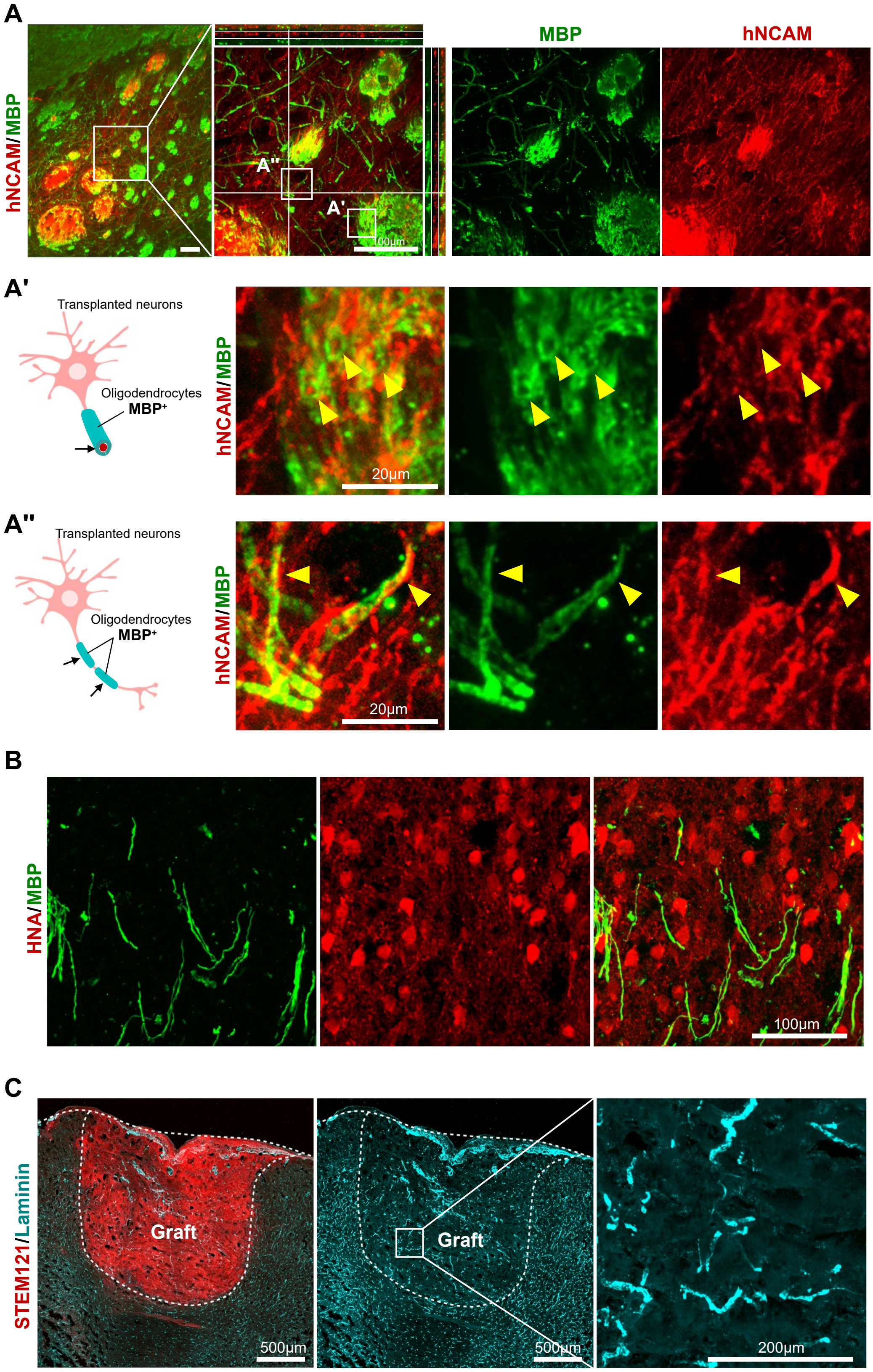


**Figure S3. Myelination and vascularization in the graft.**

(A) Immunostaining of hNCAM and oligodendrocyte marker MBP showing graft-derived axons are surrounded by mouse oligodendrocytes. The cross-sectional view is shown in panel (A'), and side view in panel (A''). Yellow arrowheads indicating the relationship between MBP and hNCAM signals. Scale bars, 100μm.

(B) Immunostaining with MBP (oligodendrocyte marker) and HNA showing no expression of MBP in transplanted cells. Scale bars, 100μm.

(C) Immunostaining with STEM121 and endothelial marker Laminin on the brain slices of 12-month after transplantation, showing the mouse blood vessel endothelium grew into the graft. Scale bars, 500μm. 200μm in magnification.

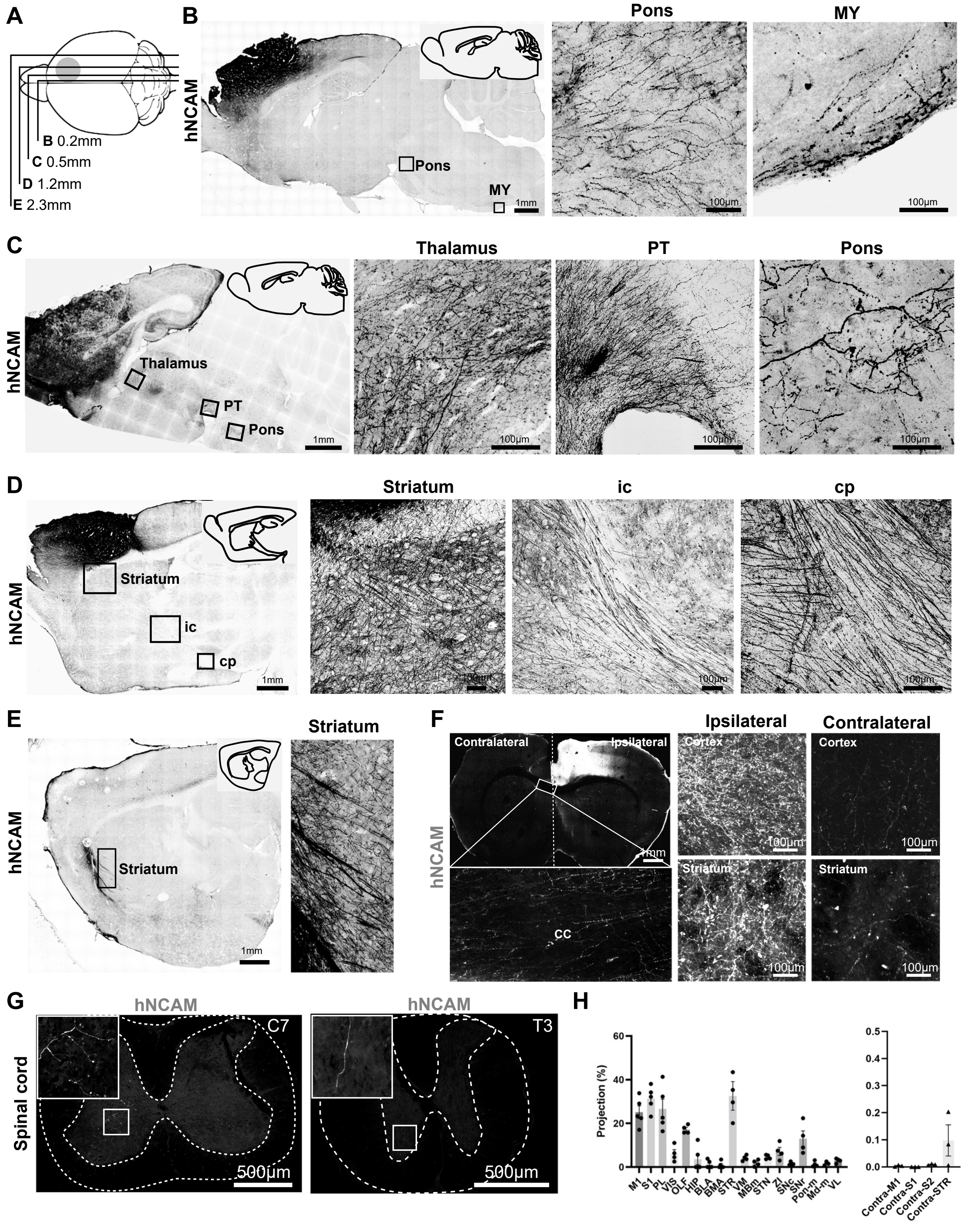


**Figure S4. Transplanted human cortical neurons rebuild the motor related neural circuits.**

(A) Schematic diagram showing the planes for brain sections.

(B to E) Immunostaining of hNCAM showing axon projections by the grafted neurons in Pons and MY (B), thalamus, PT and Pons (C), striatum, ic and cp (D), and S1 and striatum (E), in stroke mice 12 months post transplantation. Scale bars, 1mm. 100μm in magnification.

(F) Immunostaining for hNCAM showing axon projections by the grafted neurons in the contralateral and ipsilateral hemispheres at 12 months after transplantation. Scale bars, 1mm. 100μm in the magnification.

(G) Immunostaining for hNCAM showing graft-derived axons in the contralateral spinal cord C7 (top) and T3 (bottom) at 12 months after transplantation. Scale bars, 500μm.

(H) Quantification of axon projection of grafted neurons in brain regions.

All Data are presented as mean±SEM.

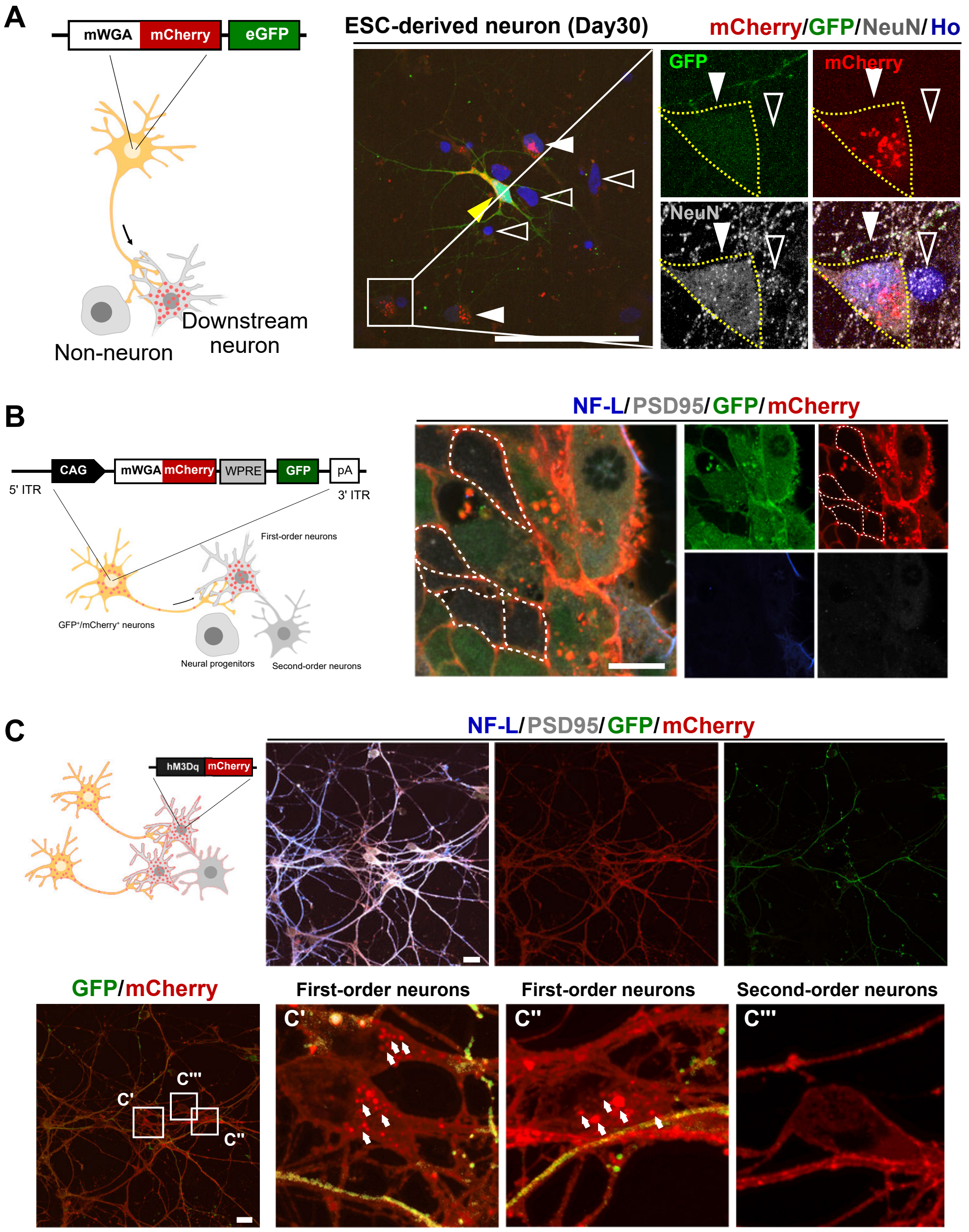


**Figure S5. Validation of anterograde mono-synaptic tracer mWmC.**

(A) Schematic of generating GFP/mWmC cell lines to validate mWmC (left). Immunostaining for NeuN, mCherry, and GFP showing that mCherry(mWmC) is only detected in NeuN+ cells (right). Yellow arrowheads indicate primary neuron (GFP+/mCherry+); White arrowheads indicate NeuN+ cells; open arrowheads indicate NeuN- cells. Scale bars, 100μm.

(B) Immunostaining of NF-L, PSD95, GFP, and mCherry on neural progenitors showing that the mWmC is not presented in GFP-/PSD95- neural progenitors. Dashed lines indicate GFP-/PSD95- cells. Scale bars, 20μm.

(C) Schematic diagram showing that primary neurons (GFP+/mWmC+) project axons distally to downstream neurons (hM3Dq-mCherry), validating that mWmC works as monosynaptic tracer. Immunostaining of NF-L, PSD95, GFP, and mCherry on downstream neurons showing no puncta red signals are detected in the second-order neurons. Arrows indicate puncta red signals in the cytoplasm. (C’ and C’’) indicate the first neurons which attach GFP+ neurites from primary neurons; C’’’ indicates the second-order neurons. Scale bars, 20μm.

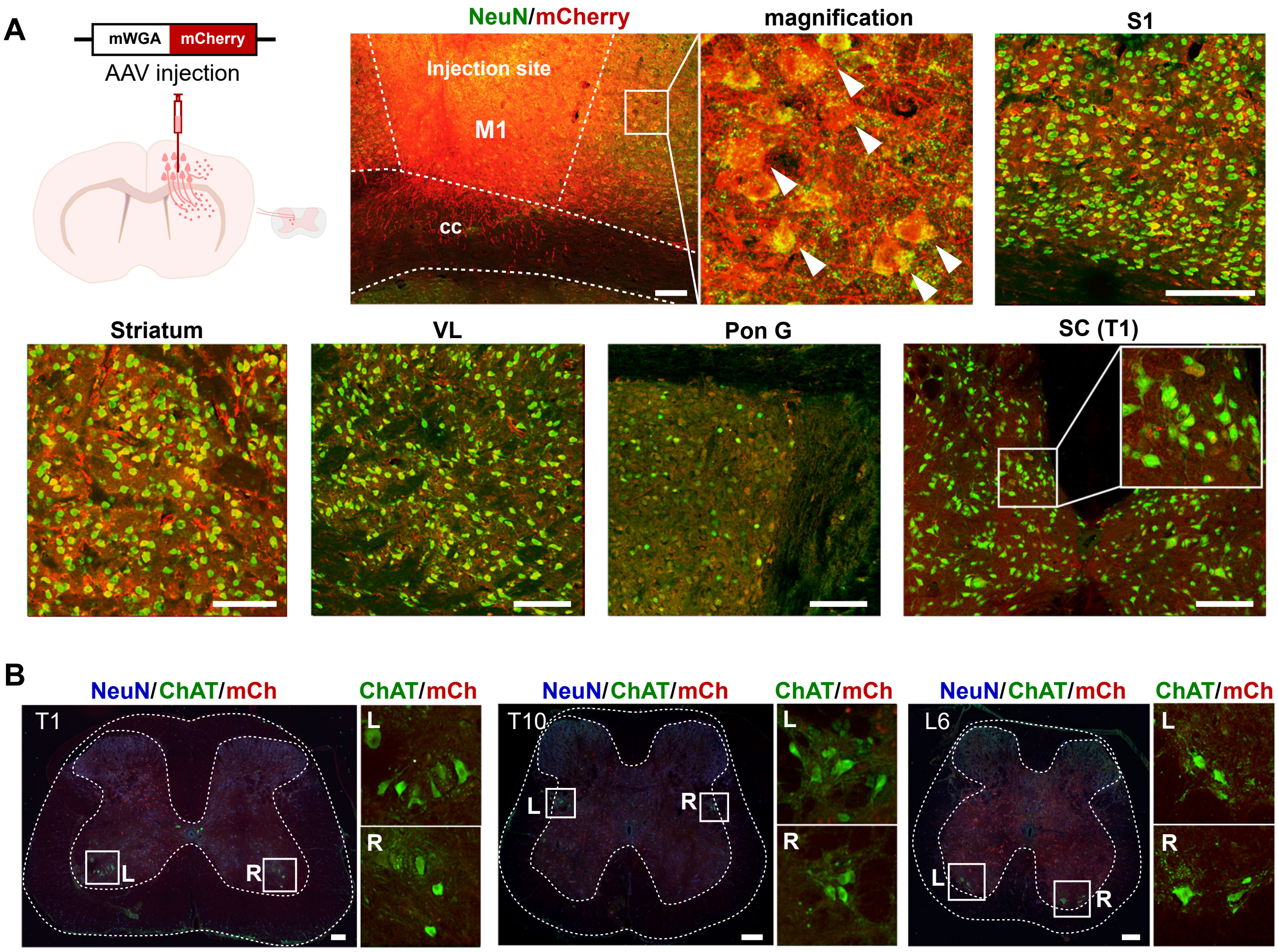


**Figure S6. Anterograde trans-synaptic tracer mWmC in vivo.**

(A) Strategy for injection of mWmC-AAV in M1c of the mouse brain. Immunostaining showing mCherry (mWmC) detection in the S1, striatum, VL, Pon G and T1 (spinal cord) in the mouse CNS. Scale bars, 200μm.

(B) Immunostaining showing mCherry (mWmC) detection in the T1, T10, and L6 (spinal cord). Scale bars, 100μm.

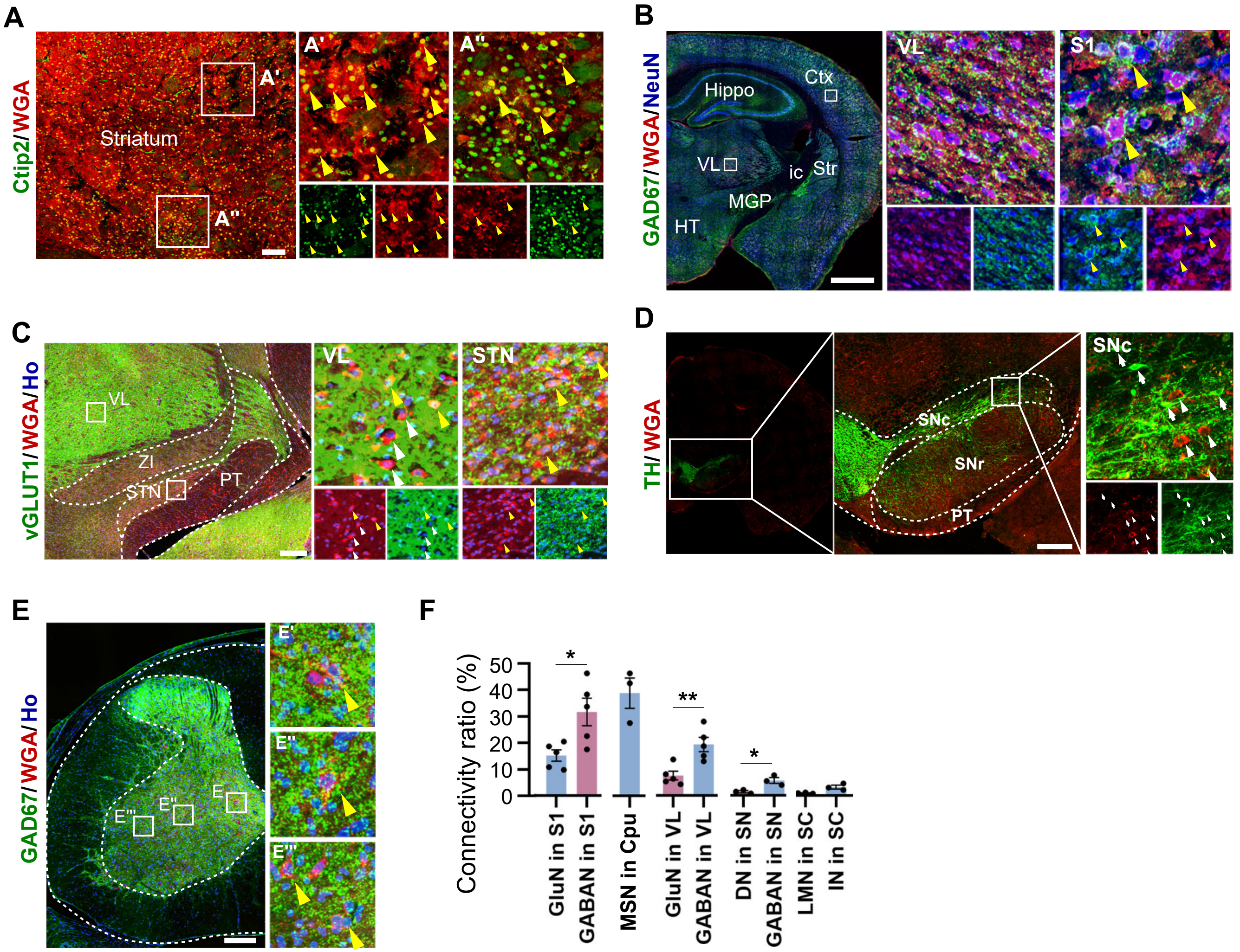


**Figure S7. Transplanted neurons synapse with specific target neurons.**

(A) Immunostaining for WGA and Ctip2 showing that host medium spiny neurons (Ctip2$^+$) were labelled by WGA in the striatum at 12-mpt. Right panels show magnified view of the insets in the dorsal (A') and ventral (A") striatum. Yellow arrowheads indicate WGA$^+$/Ctip2$^+$ neurons. Scale bars, 100μm.

(B) Immunostaining with NeuN, WGA, and GAD67 showing that GABAergic neurons in the cortex and thalamus were WGA positive. Right panels show magnification of VL and S1. Yellow arrowheads indicate WGA$^+$/GAD67$^+$/NeuN$^+$ cells. Scale bars, 1mm.

(C) Immunostaining for vGLUT1 and WGA showed that WGA labels (vGLUT1$^+$) glutamatergic neurons in the VL and STN. The magnification was shown in the right panels. Yellow arrowheads indicate WGA$^+$/vGluT1$^+$ cells. White arrowheads indicate WGA$^+$/ vGluT1$^-$ cells. Scale bars, 100μm.

(D) Immunostaining showing no overlap of WGA with hostdopaminergic neurons (TH+) in the substantia nigra. White arrowheads indicate WGA+/ TH- cells. White arrows indicate TH+/WGA- cells. Scale bars, 100μm.

(E) Immunostaining showing GAD67-expressing WGA-labeled host neurons in the spinal cord. The insets are magnified on the right, yellow arrowheads indicate WGA$^+$/ GAD67$^+$ cells. Scale bars, 100μm.

(F) Quantification of the ratio of host neurons receiving output connections of grafted cells in different brain regions. *p<0.05, **p<0.01, two-sided Student's t-test. Data are presented as mean±SEM.

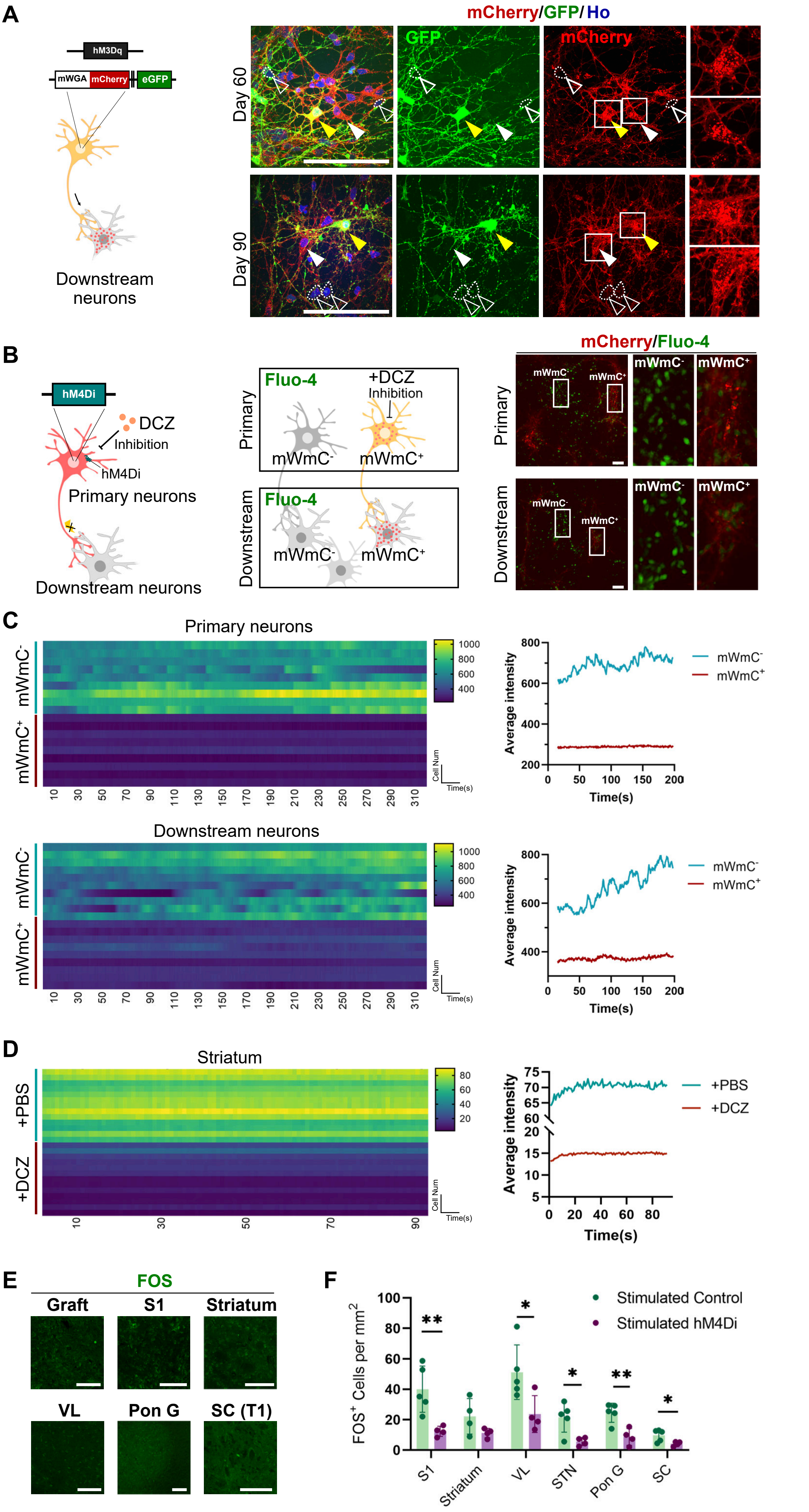


**Figure S8. Functional tests of DREADD/mWmC ESC-derived neurons.**

(A) Strategy for generation of hM3Dq-mCherry–mWmC-eGFP cell line (left). Immunostaining showing expression of GFP and mCherry(mWmC) in 2D-cultured neurons 60 (right top) and 90 (right bottom) days of induction. Scale bars, 100μm.

(B) The left panel shows the schematic of hM4Di (DREADD)/mWmC neurons following DCZ treatment (inhibition). The middle panel shows the experimental strategy. The right panel shows the calcium imaging of upstream neurons and their downstream neurons following DCZ treatment. Scale bars, 100μm.

(C) Heatmap showing the neuronal activity of mWmC+ and mWmC- neurons after DCZ addition. The top panel showing the upstream neurons. The bottom panel showing the downstream neurons. The right panel showing the average intensity over time.

(D) Heatmap showing neuronal activity of mWmC+ neurons in the striatum of mice transplanted with DREADD neurons after DCZ/PBS administration. The right panel showing the average intensity over time.

(E) Immunostaining for FOS showing the expression of FOS in host neurons in the presence of DCZ at 12-month after hM4Di neural transplantation. Scale bars, 200μm.

(F) Quantification of FOS-expressing cells in the host brain and spinal cord in stroke mice. *$p<0.05$, **$p<0.01$, Student's t-test.

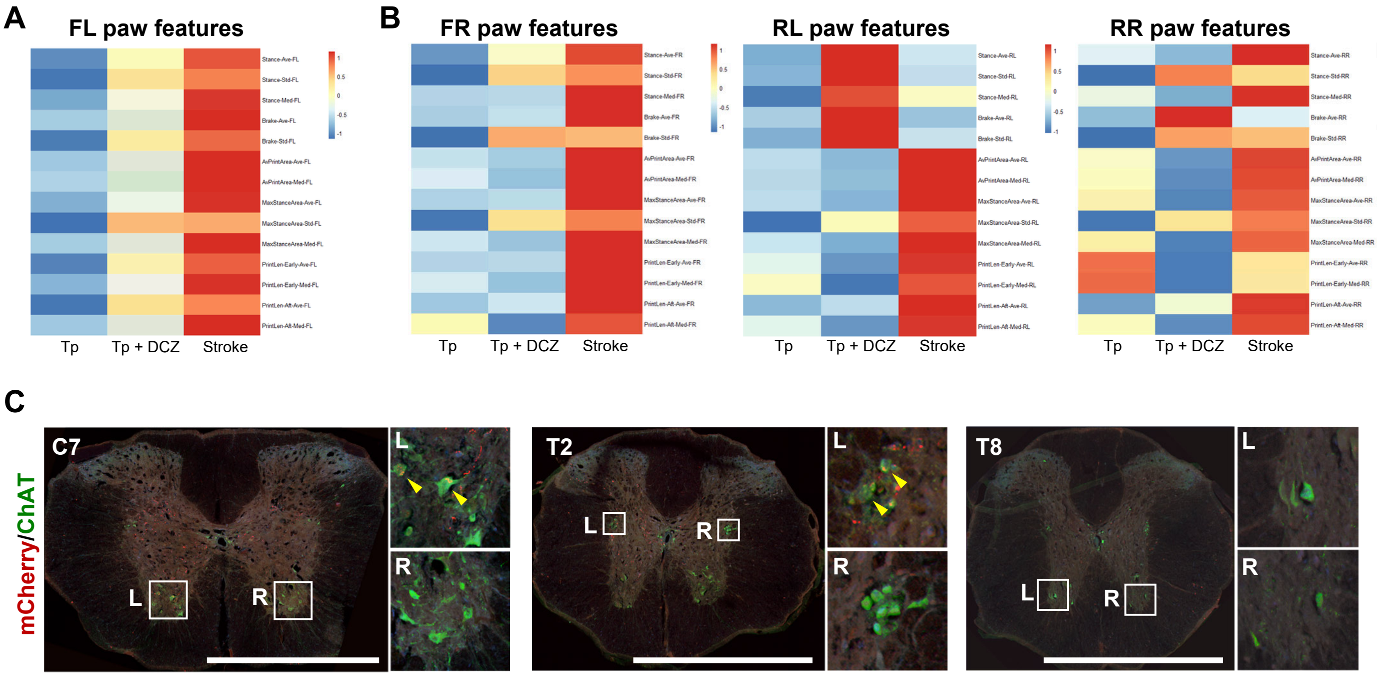


**Figure S9. Association between limb movement and spinal motor neuron connections.**

(A and B) Heatmap of parameters of front left paw (FL), front right paw (FR), rear left paw (RL), and rear right paw (RR) in treadscan assessment of mice from transplantation (Tp), transplantation with DCZ injection (Tp + DCZ; inhabitation), and Stroke group. (A) shows FL. (B) shows FR, RL, and RR.

(C) Immunostaining for mCherry and ChAT showing the synaptic connections between grafted neurons and spinal motor neurons in C7 and T2. Scale bars, 1mm.

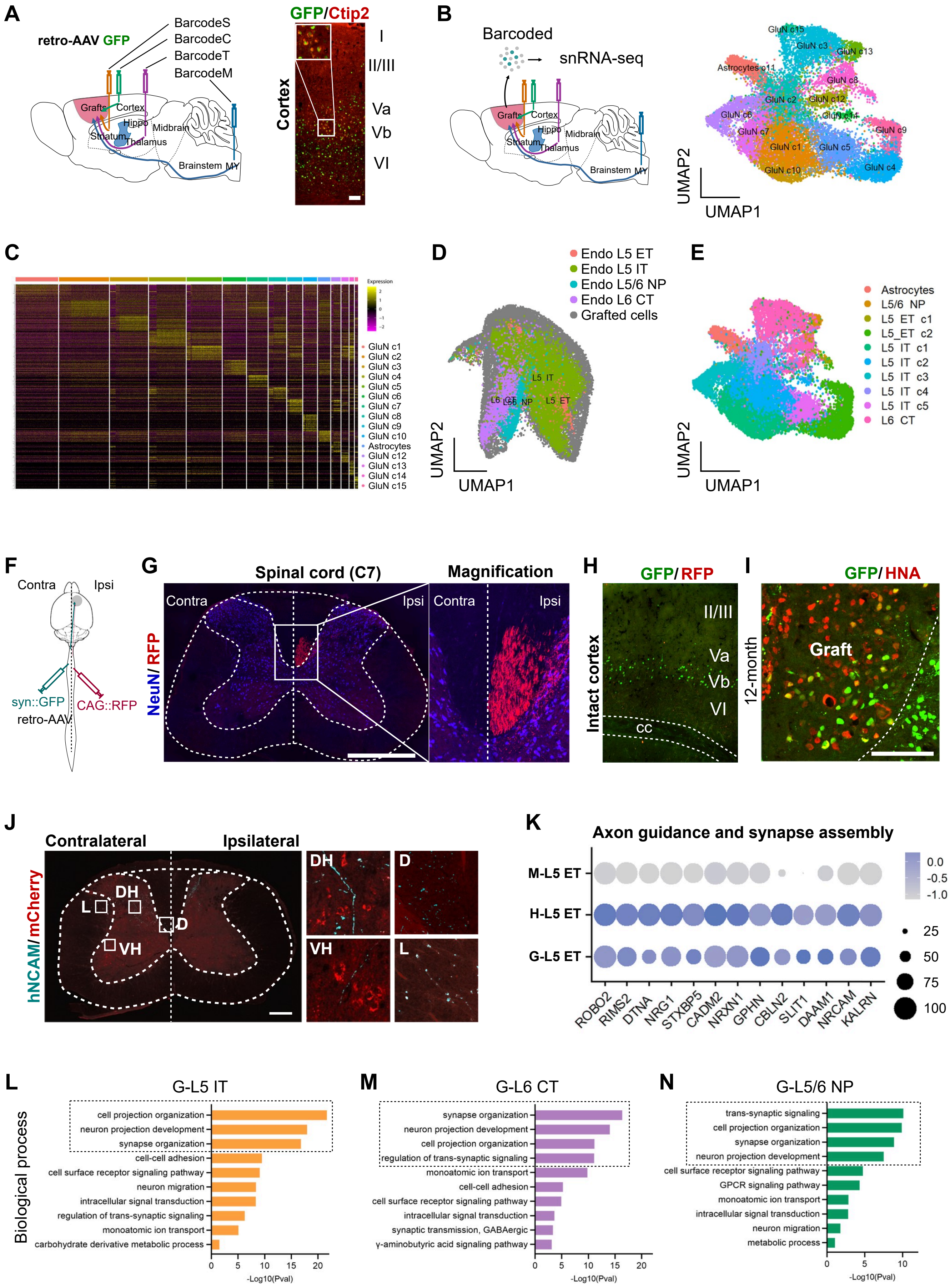


**Figure S10. Retrograde tracing for transplanted neurons.**

(A) Strategy for injection of retrograde-AAV GFP with barcodes in the transplanted mouse brain (left). Immunostaining for GFP and Ctip2 showing retrograde-AAV labelled layer V neurons in the intact cortex (right). Scale bars, 200μm.
(B) Strategy for MRT-seq at 5 weeks after AAV injection. UMAP showing clusters of nuclei from transplanted cells extracted from the mouse brain with barcoded retro-AAV-GFP injection.
(C) Heatmap showing marker expression of clusters in (B).
(D) UMAP showing the grafted cells and human M1c snRNA-seq reference dataset after Seurat integration.
(E) UMAP showing clustering of nuclei from grafted cells using label transferred from the human M1c snRNA-seq reference dataset.
(F) Strategy for retrograde AAV injection in the spinal cord of transplanted stroke mice at 12 months after transplantation.
(G) Immunostaining for RFP and NeuN showing retrograde AAV-RFP-labelled axons in the ipsilateral spinal cord of transplanted stroke mice. Scale bars, 500μm.
(H) Immunostaining of retrograde-AAV-GFP and RFP-labeled neurons in the cortex of stroke mice. Scale bars, 200μm.
(I) Immunostaining for GFP and HNA showing the transplanted neurons were labelled by retro-AAV. Scale bars, 200μm.
(J) Immunostaining showing axonal projection (hNCAM+) and output connection (mCherry+) of grafted cells to the spinal cord (C8). DH, dorsal horn. D, dorsal funiculus. VH, ventral horn. L, lateral funiculus. Scale bars, 200μm.
(K) Dot plot of top axon guidance/synapse assembly genes of human L5 ET versus mouse L5 ET neurons. The genes expression on Grafted L5 ET neurons is represented at the bottom of the plot for top feature genes.
(L-N) Bar plot of enriched Gene Oncology – Biology Process terms of cluster G-L5 IT (L), G-L6 CT (M), and G-L5/6 NP (N) in the graft dataset.

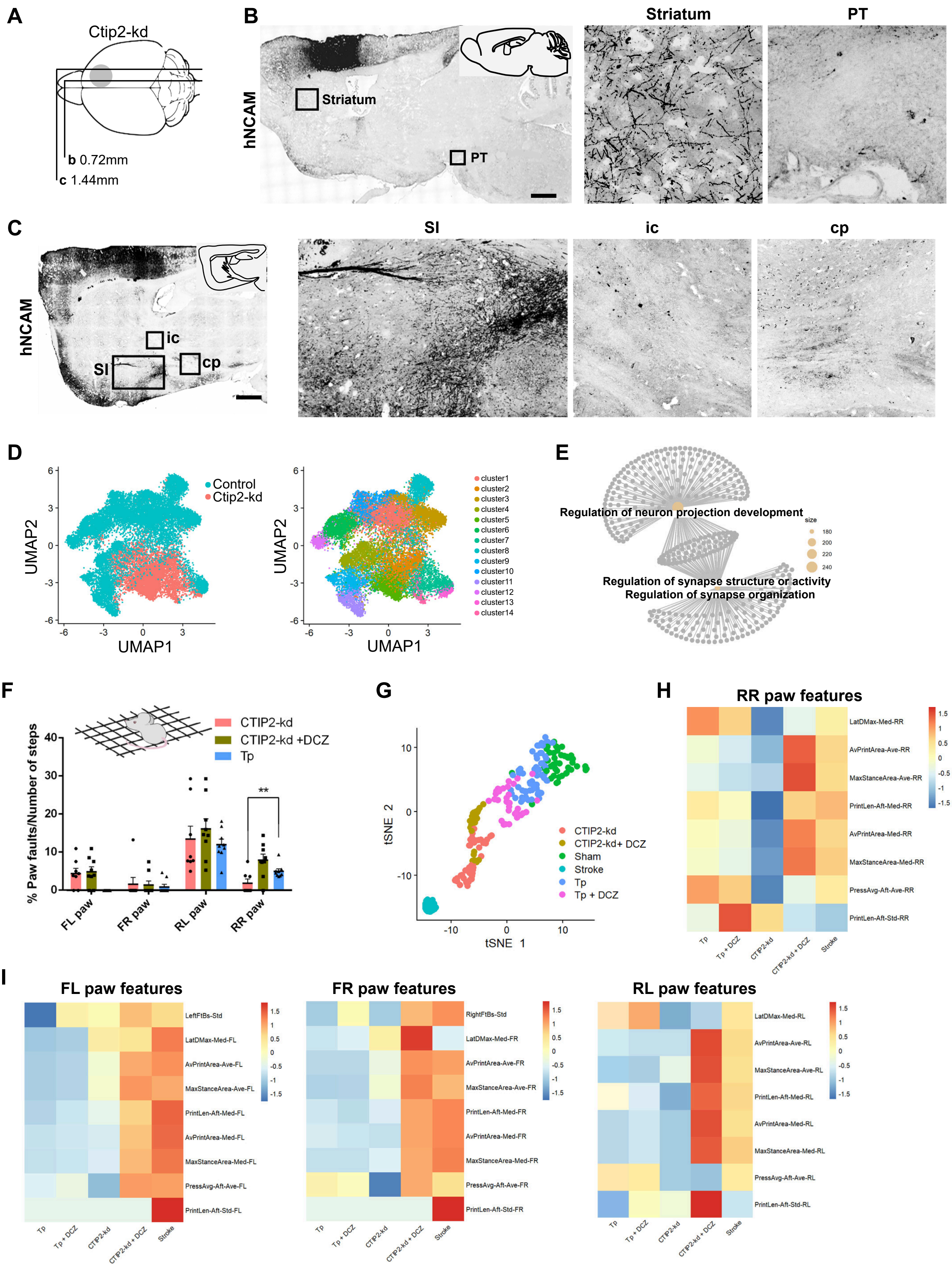


**Figure S11. The projection pattern of transplanted CTIP2-kd neurons.**

(A) Schematic for brain sections of mice transplanted with CTIP2-kd human neurons.
(B-C) Immunostaining of hNCAM showing axon projections by the grafted CTIP2-kd neurons in the striatum and PT (B), SI, ic and cp (C) at 12 months post transplantation. Scale bars, 1mm.
(D) UMAP visualization shows the subtypes of transplanted cells derived from control and CTIP2-kd cell lines after Seurat integration.
(E) Network of enriched Gene Oncology–Biology Process terms of cluster CTIP2-kd neurons versus control neurons.
(F) Barplot showing the proportion of paw faults over number of steps in grid walk assessment of mice with CTIP2-kd/hM4Di neurons transplantation, CTIP2-kd/hM4Di neurons transplantation with DCZ injection and with WT neurons transplantation.
(G) UMAP showing clustering of treadscan assessments of mice from 6 groups: Sham, Stroke, Tp (transplantation), Tp + DCZ (WT neurons transplantation with DCZ injection), CTIP2-kd (CTIP2-kd/hM4Di neurons transplantation), CTIP2-kd + DCZ (CTIP2-kd/hM4Di neurons transplantation with DCZ injection).
(H and I) Heatmap of parameters of front left paw (FL), front right paw (FR), rear left paw (RL), and rear right paw (RR) in treadscan assessment of mice from Tp, Tp + DCZ, CTIP2-kd, CTIP2-kd + DCZ and Stroke group. (H) shows FL. (I) shows FR, RL, and RR.

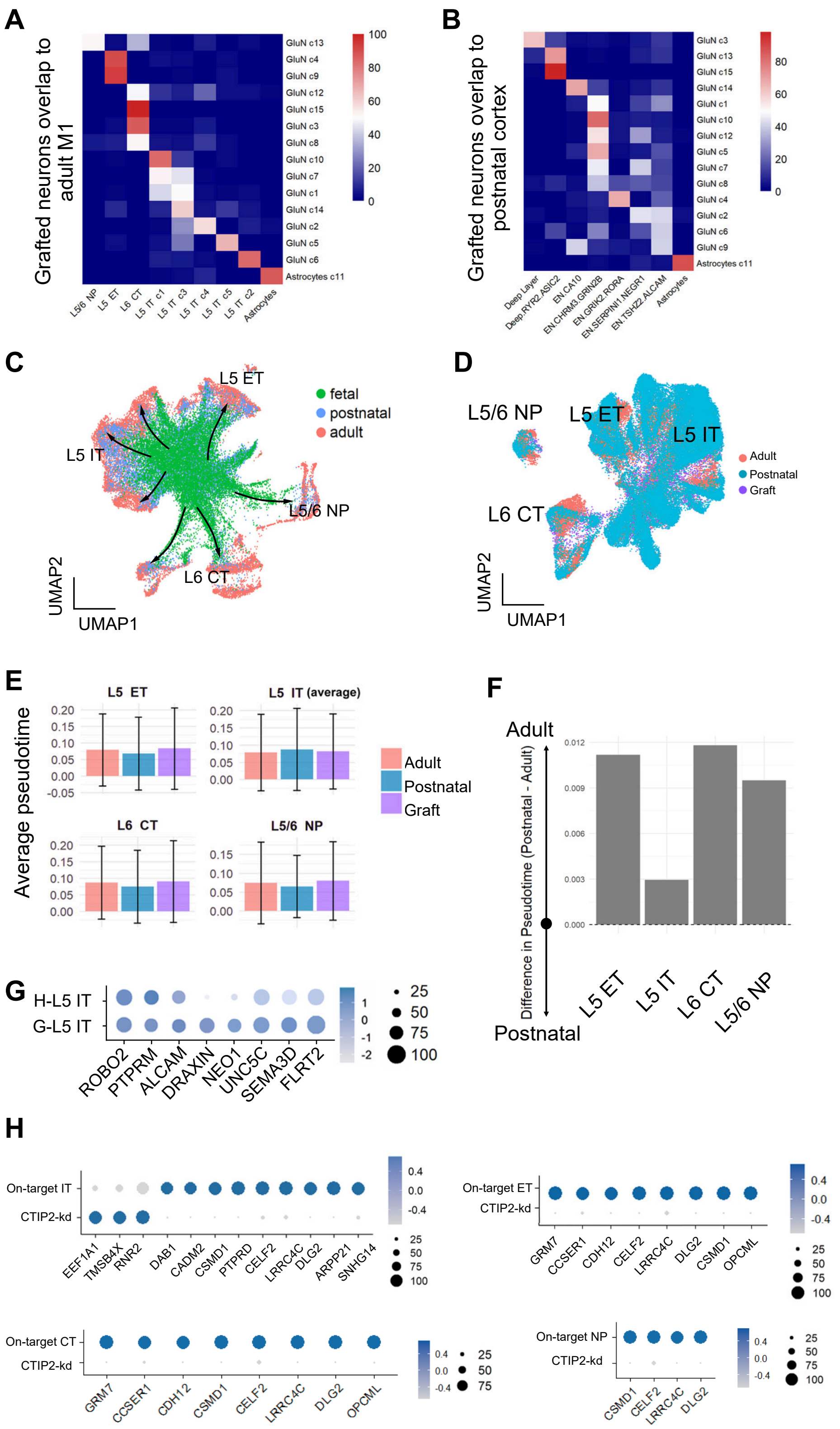


**Figure S12. Comparison of transplanted neurons with endogenous human neurons.**

(A and B) Heat maps of grafted cell clusters overlap by RNA-seq integration with human adult (A) and postnatal (B) cortical cell clusters.

(C) UMAP plots showing the maturation trajectory in each subtype of human cortical neurons from fetal, postnatal, and adult cortex.

(D) UMAP plots showing the transplanted, postnatal, and adult cortical neurons after Seurat integration.
(E) Quantification of pseudotime in each subtype of cortical neurons.
(F) Bar plot showing the difference in pseudotime distance between grafted neurons and postnatal versus adult neurons, indicating their relative similarity.
(G) Dot plot of canonical guidance gene expression on G-L5 ET neurons. The genes expression on human L5 ET neurons (H-L5 ET) is represented on the top of the plot for top feature genes.
(H) Dot plot showing the profiles of the minimal number of genes required to identify CTIP2-kd neurons from on-target grafted neurons in four subtypes.

### STAR★METHODS

### KEY RESOURCES TABLE

| REAGENT OR RESOURCE | SOURCE | IDENTIFIER |
|---|---|---|
| Antibodies | | |
| anti-Ctip2<br>1:300 | Abcam | Cat#ab18465 |
| anti-Fezf2<br>1:200 | Abcam | Cat#ab69436 |
| anti-Tbr1<br>1:1000 | Santa Cruz | Cat#sc-376258 |
| anti-Tbr1<br>1:1000 | Proteintech | Cat#20932-I-AP |
| anti-HNA<br>1:1000 | Merck Millipore | Cat#MAB1281 |
| anti-hNCAM<br>1:300 | Santa Cruz | Cat#sc-106 |
| anti-STEM121<br>1:500 | Takara | Cat#Y40410 |
| anti-Sox9<br>1:600 | R&D Systems | Cat#AF3075 |
| anti-NeuN<br>1:500 | Merck Millipore | Cat#MAB377 |
| anti-mCherry<br>1:500 | Abcam | Cat#ab167453 |
| anti-WGA<br>1:300 | Vector Labs | Cat#AS-2024-1 |
| anti-GAD67<br>1:500 | Merck Millipore | Cat#MAB5406-25UG |
| anti-vGluT1<br>1:500 | Invitrogen | Cat#9129S |
| anti-TH<br>1:500 | Immunostar | Cat#22941 |
| anti-ChAT<br>1:500 | Merck Millipore | Cat#AB144P |

| | | |
|---|---|---|
| anti-FOS<br>1:200 | Synaptic Systems | Cat#226 308 |
| anti-Iba1<br>1:500 | Wako Pure Chemical Corporation | Cat#019-19741 |
| anti-Laminin<br>1:1000 | Novusbio | Cat#NB300-144 |
| anti-MBP<br>1:500 | Abcam | Cat#ab218011 |
| Bacterial and virus strains | | |
| Lenti-CAG-mWmC-GFP | This paper | NA |
| Lenti-EF1a-mWmC | This paper | NA |
| AAV2/9-CAG-mWmC | This paper | NA |
| AAV9-syn-GCaMP | Addgene | Cat#104488-AAV9 |
| pRS-hU6-Ctip2shRNA-GFP-SV40-Puro | OriGene | Cat#TR306424 |
| Chemicals, peptides, and recombinant proteins | | |
| SB431542 | STEMCELL | Cat#72232 |
| DMH-1 | STEMCELL | Cat#100-1043 |
| CHIR99021 | Tocris | Cat#4423 |
| BDNF | Peprotech | Cat#450-02 |
| GDNF | Peprotech | Cat#450-10 |
| Compound E | MedChemExpress | Cat#HY-14176 |
| Maraviroc | MedChemExpress | Cat#HY-13004 |
| cAMP | Sigma-Aldrich | Cat#A9501 |
| Rho kinase Inhibitor | Calbiochem | Cat#555550 |
| Matrigel | Corning | Cat#VMA00048 |
| Fluoromount-G | SouthernBiotech | Cat#0100-01 |
| Hoechst | Life Technologies | Cat#33342 |
| Fluo-4, AM, cell permeant | Invitrogen | Cat#F14201 |
| Deschloroclozapine | Tocris | Cat#7193 |
| N-2 supplement (100x) | GIBCO | Cat#17502048 |
| B-27 supplement (50x) | GIBCO | Cat#17504044 |
| GlutaMAX™ supplement | GIBCO | Cat#35050061 |
| mTeSR PLUS medium | STEMCELL | Cat#100-0276 |

| | | |
|---|---|---|
| Neurobasal™ Medium | GIBCO | Cat#21103049 |
| DMEM/F-12 | GIBCO | Cat#21331020 |
| DMEM, high glucose | GIBCO | Cat#11965092 |
| TrypLE Express Enzyme (1X), no phenol red | GIBCO | Cat#12604021 |
| Dispase II, powder | GIBCO | Cat#17105041 |
| Fibrinogen | Sigma | Cat#F3879 |
| $CaCl_2$ | Sigma | Cat#499609 |
| Dimethyl Sulfoxide (DMSO) | Sigma | Cat#D8418-500ML |
| Certified Fetal Bovine Serum (FBS) | Biological Industries | Cat#04-002-1A |
| Experimental models: Cell lines | | |
| H9 hESCs | WiCell | Cat#WA09 |
| DREADD hESCs | This paper | NA |
| hM4Di hESCs | This paper | NA |
| Syn-mWmC hESCs | This paper | NA |
| DREADD/mWmC hESCs | This paper | NA |
| hM4Di/mWmC/GFP hESCs | This paper | NA |
| hM4Di/mWmC hESCs | This paper | NA |
| Experimental models: Organisms/strains | | |
| Mouse: CB17.Cg-*Prkdc*$^{scid}$*Lyst*$^{bg\text{-}J}$/Crl (SCID Beige) | Charles river | Cat# CRL:250; RRID:IMSR_CRL: 250 |
| Oligonucleotides | | |
| sgRNA: human SYN1 | This paper | NA |
| Barcode-S: GCTACGCCATCGCAAGGCCT | This paper | NA |
| Barcode-C: CCAAGCGTATGCTACGCGTT | This paper | NA |
| Barcode-T: AGCCTTGCATCGCATCAAGG | This paper | NA |
| Barcode-M: TACTGAGTCGCAAGCGCTCC | This paper | NA |
| Recombinant DNA | | |
| mWmC (shown below) | IDT | NA |
| Software and algorithms | | |
| Fiji ImageJ | Open source | https://fiji.sc |
| Graphpad Prism 9 | Graphpad | https://www.graphpad.com/ |
| Rstudio | Rstudio | https://rstudio.com |

| | | |
|---|---|---|
| Seurat 3.1.4 | Stuart et al. | https://satijalab.org/seurat |
| Imaris 10.2 | Oxford instruments | https://imaris.oxinst.com/ |
| Imaris Stitcher 10.2 | Oxford instruments | https://imaris.oxinst.com/ |

### EXPERIMENTAL MODEL AND STUDY PARTICIPANT DETAILS

All experiments were carried out according to the protocol approved by the Institutional Animal Care and Use Committee at Duke-NUS Medical School.

#### Cell Culture

Human embryonic stem cells (hESCs) were cultured in a feeder-free mTeSR PLUS medium (STEMCELL, CAT#: 100-0276) with 1 × mTeSR supplement, 1 × Non-Essential Amino Acids (NEAA; Gibco, CAT#: 11140050) and 1 × GlutaMAX Supplement (Gibco, CAT#: 35050061) in a Matrigel (Corning, CAT#: VMA00048) coated 6-well plate. hESCs were fed daily and passed every three days.

#### Photothrombotic stroke model

All animal studies were performed in accordance with the institutional animal care and use committee at Duke-NUS Medical School. Ischemic strokes in adult (10-12 weeks) male SCID mice were induced through photo-thrombosis. Briefly, Rose Bengal was administered intravenously at a dose of 0.1 mg/g per mouse. Mouse skull was exposed under 3% isoflurane anesthesia. The right motor cortex (anterior-posterior [AP] = +2 mm, lateral [L] = +1 mm) received a 2.5-mm diameter illumination of cold light through the intact skull for 15 min. Mice with speeds ranging from 10–12 m/s in TreadScan test were selected for inducing stroke model.

### METHOD DETAILS

#### Cell Culture

Induction of forebrain glutamate neurons was described[41]. In brief, hESCs were cultured on Matrigel-coated 6-well plates for five days, growing to a density of 80% confluency. On day 0, ESC colonies were gently detached using a 1 mL pipette to form cell aggregates. Cell aggregates were suspended in T75 flasks (Greiner Bio-One, CAT#: 658195) for 7 days with 8 mL neural induction medium (NIM) consisting of DMEM/F12 (Gibco, CAT#: 21331020), 1 × N2 supplement (Gibco, CAT#: 17502048), 1 × NEAA, 2-μM SB431542 (STEMCELL, CAT#: 72232), and 2-μM DMH-1 (STEMCELL, CAT#: 100-1043). Cell aggregates were adhered to 6-well plates in the presence of NIM with 5% FBS for 6 h, and then replaced with fresh NIM. The aggregates were cultured in NIM till neural rosette formation at day-14. The rosettes were gently blown off using 1 mL pipettes and suspended in flasks with NIM for 6 days. Then, NIM was changed every five days from day 20. The neural progenitors were maintained in NIM till transplantation or immunostaining. Progenitors were digested into single cells using TrypLE for 3 min at day 33. After two additional days of incubation in NIM supplemented with 1 × B-27 (Gibco, CAT#: 17504044) and 100 nM compound E (MedChemExpress, CAT#: HY-14176), the progenitors were collected for transplantation. For immunostaining, progenitors were seeded on glass coverslips and staining was performed after one-week culture.

### Cell Transplantation

Medium for NPC transplantation contains fibrinogen and maraviroc, as described previously[17]. Stock solution of 30 mg/mL fibrinogen, 50 mg/mL maraviroc and 250 mM CaCl2 (100×) were prepared in the following manner: fibrinogen (F3879, Sigma) was dissolved in a-CSF for 1 hour at room temperature. Maraviroc was dissolved in Dimethyl Sulfoxide (DMSO) (D8418-500ML, Sigma). $CaCl_2$ (499609, Sigma) was dissolved in deionized (DI) water. All solutions were sterile filtered and stored at -20 ℃ for use. Fibrinogen stock solution was diluted to 10 mg/mL fibrinogen in aCSF before preparing the cocktail. Cocktail was made of 30 mg/mL maraviroc and 10 mg/mL fibrinogen in a volume ratio of 1: 9 with 2.5 mM $CaCl_2$. Animals were randomly grouped and transplanted with forebrain glutamatergic progenitors. Fifty thousand cells were resuspended in 1 μL medium and injected into the injured site ([AP] = +2 mm, [L] = +1 mm, vertical [V] = -1.5 mm, from dura).

### Tissue Preparation and Immunohistochemistry

Animals were sacrificed with a lethal dose of pentobarbital (250 mg/kg) and immediately perfused with PBS followed by 4% cold paraformaldehyde (PFA). The brain samples were fixed in cold PFA overnight and immersed sequentially in 20% and 30% sucrose at 4 ℃ until sunk. For immunostaining, sections were incubated with blocking solution containing 10% normal donkey serum and 0.2% TritonX-100 for 1h at room temperature. Then sections were incubated with primary antibodies overnight at 4 ℃.

Sections were subsequently rinsed with PBS and incubated with corresponding secondary antibodies for 1h at room temperature, followed by washing with PBS for 5min × 3. Immunolabeled sections were mounted by Fluoromount-G with Hoechst.

### Whole-mount Staining, Imaging and 3D Reconstruction

The protocol of whole-mount staining has been described (Renier, 2014). Briefly, transplanted mice were perfused with 20mL PBS and 20mL 4% PFA/PBS. The dissected brains and spinal cords were fixed in 4%PFA/PBS at 4℃ overnight and 1hour RT with shaking, followed by wash in PBS with shaking for 30min for 3 times. Samples were dehydrated with 20%, 40%, 60%, 80%, and 100% methanol/$H_2O$ for 1 hour each, re-washed with 100% methanol for 1 hour and chilled at 4℃. The samples were then incubated in 66% (dichloromethane) DCM / 33% methanol at RT overnight, washed twice in 100% methanol at RT and chilled at 4℃. The samples were then bleached in chilled fresh 5% $H_2O_2$ (1 volume 30% $H_2O_2$ to 5 volumes MeOH) at 4℃ overnight and rehydrated with 80%, 60%, 40%, 20% methanol/$H_2O$ and PBS for 1 hour each at RT. The samples were washed in PBS/0.2% Triton X-100 for 1 hour twice at RT and incubated in Permeabilization Solution (PBS/0.2% Triton X-100/20% DMSO/0.3 M glycine) and then Blocking Solution (PBS/0.2% Triton X-100/10% DMSO/6% Donkey Serum) at 37℃ for 2 days each. PTwH was prepared as PBS/0.2% Tween-20 with 10 μg/ml heparin. Then the samples were incubated with primary antibody in PTwH/5% DMSO/3% Donkey Serum at 37℃ for 7 days and washed in PTwH for 4-5 times until the second day. Then the samples were incubated with secondary antibody in PTwH/3% Donkey Serum at 37℃ for 7 days and washed in PTwH for 4-5 times until the second day. After immunolabeling, the samples were dehydrated with steps described above and incubated in 66% DCM / 33% Methanol at RT with shaking for 3 hours. Then the samples were incubated in 100% DCM for 15 min with shaking twice and incubated in DBE until imaging.

Single plane illuminated (light-sheet) image stacks were acquired using the Ultramicroscope Blaze with the following filter sets: ex 470 nm, em 525/50 nm; ex 561 nm, em 595/40 nm; ex 640 nm, em 680/30 nm; ex 785 nm, em 845/55 nm (Miltenyi Biotec). The (sample) was imaged using a 4.0x/12.0x objective lens with a magnification of (insert magnification here, the available magnification is 0.66, 1, 1.67, 2.5).

Raw image data in the form of OME-TIFFs were converted into imaris format using the Imaris File Converter version (insert version), then stitched together using the Imaris Stitcher version (insert version), and analysed using the Imaris analysis software version (insert version)

The whole-mount imaging was taken using 3D light-sheet microscopy and raw data were converted into iMaris format from Miltenyi. The imaris images were 3D reconstructed with imaris stitcher 10.2.0 (RRID:SCR_007370) for final viewing.

**Retrograde Labeling**

To retrogradely label the grafted neurons associated with motor control in the SC (those projecting through SC level C7), an AAV injection was performed at 5-week before collecting grafts. Specifically, mice were anesthetized and a laminectomy of the C7-T1 vertebral processes was performed. AAV-Retro (AAV-CAG-hChR2-H134R-tdTomato) was injected at one site per segment within the ipsilateral (right side) C7-T1 spinal segments (300 nL/injection) at 0.9 μm below the dorsal spinal cord surface at each site (rate = 100 nL/min). In the same surgical session, AAV-Retro (pAAV-hSyn-HI-GFP-WPRE-SV40) was injected into the corresponding contralateral site, and 300 nl virus was injected at same depth. A total of three animals were sacrificed for snRNA sequencing.

For multiple-site retrograde tracing, 20 bp barcode sequence for identity was inserted before SV40 polyA in pAAV-hSyn-HI-GFP-WPRE-SV40 (OBiO Technology).

Barcode-S for striatal identity: GCTACGCCATCGCAAGGCCT

Barcode-C for cortex (S1Tr) identity: CCAAGCGTATGCTACGCGTT

Barcode-T for thalamus identity: AGCCTTGCATCGCATCAAGG

Barcode-M for MY identity: TACTGAGTCGCAAGCGCTCC

Mice were anesthetized with 3% isoflurane. In the same surgical session, the barcode-AAV was injected (500 nL/injection, rate = 100 nL/min) at the following coordinates from Bregma: striatum (AP = 0.26 mm, ML = 2.00 mm, DV = -2.80 mm), cortex (AP = -1.46 mm, ML = 1.50 mm, DV = -1.20 mm), thalamus (AP = -0.95 mm, ML = 1.10 mm, DV = -3.50 mm), SC (AP = -8.30 mm, ML = 0.00 mm, DV = -5.20 mm). The micropipette was left in tissue for 10 min before slowly withdrawn. A total of three mice were perfused by cold PBS, and the grafts were dissociated to single nuclei for snRNA-seq. The brain and SC were collected for immunostaining.

**snRNA Sequencing Library Preparation and Sequencing**

The single nuclei RNA-seq libraries were constructed using GEXSCOPE™ Single Nuclei RNAseq Library Kit (Singleron Biotechnologies) according to the manufacturer´s instructions. Briefly, for each library, the nuclei suspension of specified concentration was loaded onto a microfluidic chip for capture. The single-nuclei capture, lysis and mRNA capture steps were automated using Singleron Matrix NEOTM system. The final single-cell RNA sequencing libraries were sequenced on Illumina NovaSeqX plus 25B flowcell with paired-end 150 bp.

**Transcriptome Data Pre-processing**

Fastq files were preprocessed using the CeleScope® tools (version 2.0.7); www.github.com/singleron-RD/CeleScope, Singleron Biotechnologies), to generate raw data using default parameters. Briefly, cellular barcodes in Read 1 were used to demultiplex and identify reads of the same cell origin. Low quality and adapter sequences were removed using cutadapt (version 3.7). The mapping was done using STARSOLO (https://github.com/alexdobin/STAR/blob/master/docs/STARsolo.md) against human genome build (GRCh38) with ENSEMBL Gene Annotation (Version 99) with gene sequences of GFPCRE1, GFPCRE2, GFPCRE3, GFPCRE4, and CHR2RFP included. The reads were assigned to genes using the featureCount tool (https://subread.sourceforge.net) and the cell calling was performed by fitting a negative bimodal distribution and determining the threshold between empty wells and cell-associated wells. The gene count matrix was then generated, providing the number of unique molecular identifier (UMI) for each gene and cell. https://cutadapt.readthedocs.io/en/stable/installation.html).

**Single-nuclei Data Analysis**

Downstream analysis was done using the snRNA analysis pipeline CeleScoot 1.2.2 (https://github.com/singleron-RD/scrna). QC metrics, such as the number of genes detected per cell (nFeature_RNA) and the percentage of mitochondrial UMI (percent_mt) were extracted from the gene count matrix with the function calculate_qc_metrics. Nuclei with high percentage of mitochondrial counts (>20%) were filtered out. The cells with a high number of detected genes (>5000) were excluded to remove possible doublets. Cell debris characterized by low detected genes (<600) was also excluded from the data analysis. DEGs were filtered with standard threshold of log2FC > 0.25 or < -0.25 and p-value < 0.05. Projection codes for each subtype were calculated by machine learning model, full code is available at the Dryad with accession number 4j0zpc8qd (Dataset DOI: 10.5061/dryad.4j0zpc8qd).

All subsequent analysis was performed using R (version 4.4.0) package Seurat (version 5.0.3). The raw count matrices were normalized by natural-log transformation with a scale factor of 10000. Dimensionality reduction using principal component analysis (PCA) was performed, and clusters of nuclei were identified in PCA space by shared nearest-neighbour graph construction and modularity detection implemented by the FindNeighbors and FindClusters functions using a dataset dimension of 30 and resolution of 1.0. The dataset was embedded for visualization with UMAP. Marker genes for each cluster were calculated using FindMarkers with default parameters based on normalized data, and filtered with the criteria: p-value < 0.05, |log2FC| > 1 and percentage of ident.1 >

10%. We identified cell types by the expression of known marker genes. Glutamatergic neurons were identified by the expression of SYT1 and SATB2. Deep layer neurons were identified by co-expression of CTIP2 (BCL11B) and FEZF2 or TBR1. Upper layer neurons were identified by the expression of Brn2 (POU3F2). GABAergic neurons were identified by the expression of GAD1. Astrocytes were identified by the expression of AQP4 and SLC1A3. Oligodendrocytes were identified by the expression of OLIG1 and OLIG2. Proliferating cells were identified by KI67 (MKI67). To better understand the cell population, we integrated the dataset to the human adult M1 cortex dataset using the Anchor-based CCA integration in Seurat. Only genes detected in both datasets were included in comparison. Our cells were defined using the label from the reference clusters where they were overlapped. Heatmaps were drawn with 50 cells randomly selected from each cluster. Gene Ontology (GO) enrichment analysis was performed using enrichR.

**Trajectory analysis**

We performed UMAP dimension reduction at the trajectory level with the integration function from Seurat, using all genes. Each cell was assigned a pseudotime value based on its position along the trajectory using monocle3/v1.3.7 function “order_cells()”[42].For the integrated endogenous dataset, each cell was assigned a pseudotime value along the trajectory within the whole dataset. For the integrated endogenous-graft dataset, cells were subset to each subtype and each cell was assigned a pseudotime value based on its position along the trajectory within the certain subtype.

**Identification of transcriptional codes by machine learning**

Single-cell RNA sequencing data was processed and analyzed using Python. Gene expression data was extracted from the AnnData object, with raw counts converted into a dense array format. Cells were classified based on their source, distinguishing grafted cells from endogenous cells. Canonical correlation analysis (CCA) embeddings, stored in `obsm['X_cca']`, were concatenated with gene expression features to create a combined feature matrix.

Feature selection was performed in two steps. First, the top 100 features (genes) were calculated using the `SelectKBest` method with the `f_classif` statistical test. Subsequently, L1-regularized logistic regression (Lasso) was applied to further refine the feature set, retaining only genes with nonzero coefficients.

The refined feature set was used to train a logistic regression model with L1 regularization (C=0.01) on 80% of the data, while the remaining 20% was used for testing. The model was trained using the `liblinear` solver. The results were cross-validated by Bayesian Additive regression trees. Performance was assessed using receiver operating characteristic (ROC) curve analysis, and the area under the curve (AUC) was computed to evaluate classification accuracy.

**Neuron Prediction and Output**

The trained logistic regression model was applied to the full dataset to predict cell source probabilities. Cells with a probability greater than 0.5 were classified as grafted. The above pipeline was applied iteratively to multiple cell subtypes, with subtypes identified based on the `brief_id` field. The analysis was conducted separately for each subtype, ensuring subtype-specific feature selection, model training, and visualization.

**CTIP2-kd ESC Line**

Human Ctip2 (BCL11B) 29mer shRNA plasmids (pRS-hU6-Ctip2shRNA-GFP-SV40-Puro) and non-effective 29-mer scrambled shRNA cassette in pRS Vector were obtained from OriGene (CAT#: TR306424). The lentivirus was generated in the HEK 293FT cell line by transfecting with the packaging and backbone plasmids. HEK 293FT cells were cultured in DMEM high glucose (Cytiva CAT#: SH30081.01) with 10% FBS. The supernatant was collected after 3-day culture. Viral particles were concentrated by ultracentrifugation at 25 000 rpm for 2.5 h at 4 °C. The viral particles were resuspended in DMEM. The H9 ESCs were infected with the lentivirus (MOI = 50) overnight at 37 °C. The infected ESCs were sparsely cultured in the 60-mm dish (D8054, Sigma) for 3 days. The GFP labeled clones were selected for culture. The cell line was identified by staining for Ctip2 as above described.

### Anterograde tracing

mWGA-mCherry. Codon-optimized WGA (mWGA) was synthesized at IDT and fused with RFPs. We used mCherry to tag the C-terminus of mWGA. These vectors are available at Addgene for requests. The reference sequence for mWmC cDNA was optimized as below[18]:

ATGGAGACCGACACCCTGCTGCTGTGGGTGCTTCTGCTGTGGGTCCCTGGCAGCA
CTGGCGATGGGCCTGTGATGACCGCCCAAGCTCAGAGGTGCGGCGAGCAAGGCA
GCAACATGGAGTGCCCTAATAACTTGTGCTGCTCTCAGTACGGCTATTGCGGCATG
GGTGGCGACTACTGCGGCAAGGGCTGTCAGAACGGCGCCTGCTGGACTAGCAAG
AGGTGCGGCTCCCAAGCCGGCGGTGCCACCTGCCCTAACAATCACTGTTGCTCAC
AGTACGGTCACTGCGGCTTCGGCGCCGAGTACTGTGGGGCTGGTTGCCAAGGCG
GCCCTTGTAGGGCCGATATCAAGTGCGGCAGTCAAAGCGGCGGCAAATTGTGCCC
TAACAACCTGTGCTGCTCTCAGTGGGGTTTCTGCGGACTGGGAAGCGAGTTTTGC
GGCGGCGGGTGTCAATCCGGCGCTTGTAGCACCGACAAGCCTTGCGGCAAGGAC
GCCGGCGGAAGGGTGTGCACCAACAACTACTGCTGCAGCAAATGGGGATCGTGT
GGCATAGGCCCTGGCTACTGCGGCGCTGGGTGTCAGTCGGGCGGCTGCGACGCC
GCTAGGGACCCTCCTGTGGCAAGCGCCACCATGGTGAGCAAGGGCGAGGAGGAC
AACATGGCCATCATCAAGGAGTTCATGAGGTTCAAGGTGCACATGGAGGGCAGCG
TGAACGGCCACGAGTTCGAGATCGAGGGCGAGGGCGAGGGAAGGCCTTACGAGG
GCACACAGACCGCCAAGCTGAAGGTGACCAAGGGCGGCCCTCTGCCTTTCGCCT
GGGACATCCTGAGCCCTCAGTTCATGTACGGCAGCAAGGCCTACGTGAAGCACCC
TGCCGACATCCCTGACTACCTGAAGCTGAGCTTCCCTGAGGGCTTCAAGTGGGAG
AGGGTGATGAACTTCGAGGACGGCGGCGTGGTGACCGTGACCCAAGACAGCAGC
CTGCAAGACGGCGAGTTCATCTACAAGGTGAAGCTGAGGGGCACCAACTTCCCTA
GCGACGGCCCTGTGATGCAGAAGAAGACCATGGGCTGGGAGGCAAGCAGCGAGA
GGATGTACCCTGAGGACGGCGCCCTGAAGGGCGAGATCAAGCAGAGGCTGAAGC
TGAAGGACGGCGGCCACTACGACGCCGAGGTGAAGACCACCTACAAGGCCAAGA
AGCCTGTGCAGCTGCCTGGCGCCTACAACGTGAACATCAAGCTGGACATCACAAG

CCACAACGAGGACTACACCATCGTGGAGCAGTACGAGAGGGCCGAGGGAAGGCA
CAGCACCGGCGGCATGGACGAGCTGTACAAGTGA

For adult motor cortex tracing, animals were anesthetized with isoflurane and stereotaxically injected (primary motor cortex: AP = 0.85mm, ML = 1.50 mm, DV = -2.00 mm) with AAV-CAG-mWmC (OBiO Technology). The viruses, animal lines and injection sites used are as follows. For Figure S4C, AAV-CAG-mWmC (500 nl) (100 nl/ min) was injected into the NSG mice. A total of five mice were sacrificed for immunostaining at four weeks after AAV injection.

To generate the SYN1-mWmC hESC line, the AAVS1-puro-hSyn-mWmC construct was integrated into the AAVS1 locus of H9 hESCs using the CRISPR/Cas9 system (TrueCut™ Cas9 Protein v2, A36497, Life Technologies) via transfection with the NEON 10 µL Transfection Kit (MPK1025, Life Technologies).

The single guide RNA (sgRNA) sequence used for targeting the AAVS1 locus was as follows: 5`-
mU*mG*mU*rCrCrCrUrArGrUrGrGrCrCrCrArCrUrGrGrUrUrUrArGrArGrCrUrAr
GrArArArUrArGrCrArArGrUrUrArArArArUrArArGrGrCrUrArGrUrCrCrGrUrUrArUrCrArAr
CrUrUrGrArArArArArGrUrGrGrCrArCrCrGrArGrUrCrGrGrUrGrCmU*mU*mU*rU-3`

The DREADD hESC line was described previously[43]. To generate the DREADD-CAG-mWmC and DREADD-CAG-mWmC-GFP hESC lines, the DREADD hESC line was infected with the lentivirus. Specifically, the mWmC cDNA was inserted into plasmid lenti-CAG-WPRE or lenti-CAG-IRES-GFP-WPRE. The lenti-CAG-mWmC-GFP and lenti-CAG-mWmC virus were generated as described in “Ctip2-kd ESC Line”. The DREADD hESCs were infected with the lentivirus (MOI = 100) overnight at 37 °C. The infected ESCs were sparsely cultured in the 60-mm dish. After three-day culture, six clones for each line were collected separately into a 24-well plate. The hESCs were cultured on coverslips for WGA staining to identify the cell line.

**Imaging and Cellular Quantification**

Serial coronal (1.54 mm to -0.22 mm from Bregma) sections were collected on a freezing microtome at a 30-μm thickness and stored at -20 ℃. To quantify the cell number of Ctip2, Fezf2, Tbr1, HNA, Sox9, WGA, mCherry, GFP, FOS, Hoechst, vGluT1, GAD67, TH, ChAT, and NeuN positive cells, one brain slice was selected from every 6 serial slices for stereological counting on a Zeiss M1 Microscope with Stereo Investigator software (MBF Bioscience). In brief, immunolabeled slices were scanned on Zeiss M1 Microscope, grafts area was outlined manually, and then corresponding fluorescence labeled cells were unbiasedly counted. Data were replicated 4 to 5 times per group. All data were expressed as means±SEM.

To quantify the population of Ctip2, Fezf2, and Tbr1 expressing cells among total $HNA^+$ cells on 16mm coverslips (Marienfeld Superior), all coverslips were scanned and captured by 20× objective with a confocal microscope (Nikon), and then was counted with the ImageJ software. Data were replicated three times. All data were expressed as means±SEM.

All fluorescence images of sections and coverslips were scanned and acquired using a confocal microscope (Nikon) with a 20×, 40×, or 100× objective. Live calcium imaging of cultured neurons and brain slices was recorded using a confocal microscope (Nikon) with a 20× objective.

**Behavioral Tests**

Before inducing stroke models, mice were screened using NeurodegenScan Suite (Clever Sys Inc.) for gait analysis. The mice were placed on a stationary treadmill, which accelerated to 8 m/s within 5 seconds. Mice with speeds ranging from 10–12 m/s were selected for inducing stroke model.

Behavioral tests were conducted 12 months after transplantation. During these tests, mice were placed on a stationary treadmill that accelerated to 8 m/s within 5 seconds. One week prior to the formal test, the mice underwent training sessions every two days. Each training session consisted of three practices, with 30 minutes between each practice. For each animal, a 20-second video was recorded for data analysis, with 20–25 animals per group. Each animal was tested three times for subsequent analysis using nonlinear dimensionality reduction. GaitScan Analysing System (TreadScan v.4) outputs the detailed results of these parameters from analysis into Microsoft Excel files and gives statistical results.

For DCZ administration, mice transplanted with hM4Di-neurons underwent behavioral testing. One hour after the initial test, the mice received an intraperitoneal injection of DCZ. Behavioral testing was repeated 40 minutes after the DCZ administration.

### QUANTIFICATION AND STATISTICAL ANALYSIS

A two-tailed Student's t-test was used for the pairwise comparison between two groups. The rest of the data were analyzed using a one-way or two-way ANOVA as appropriate. Error bars in all figures represent means ± SEM. Differences were considered statistically significant at a P value of less than 0.05. All data were analyzed using GraphPad Prism.
